# CNTD1 plays crucial roles in prophase I progression and crossover designation during female meiosis and is critical for establishing the ovarian reserve

**DOI:** 10.1101/2023.09.25.559118

**Authors:** Anna J. Wood, Rania M. Ahmed, Leah E. Simon, Rachel A. Bradley, Ian D. Wolff, Paula E. Cohen

## Abstract

In meiotic prophase I, hundreds of double-strand breaks (DSBs) are formed throughout the genome. A majority of these breaks are repaired as non-crossovers (NCOs), while a minor subset are repaired as crossovers (CO). COs are essential for the faithful segregation of homologous chromsomes at the end of prophase I and errors in CO designation can result in aneuploidy, germ cell death, birth defects, or infertility. These errors are more evident in female meiosis compared to males and suggests that the events of meiotic prophase I are sexually dimorphic with respect to CO formation, placement, resolution, and/or surveillance. Here, we demonstrate a critical role for Cyclin N-Terminal Domain Containing 1 (CNTD1) protein in ensuring appropriate CO frequency and distribution across the genome during meiosis in females. We find that CNTD1 localizes with the heterodimer, MutLγ, which marks the majority of CO that emerge in pachynema of prophase I, implicating CNTD1 in late-stage CO designation and/or maturation. Accordingly, loss of *Cntd1* in oocytes results in failure to load MutLγ and thus results in a catastrophic loss of chiasmata and sterility. Further investigation yielded a distinct phenotype in which the primordial follicles that form upon dictyate arrest are steadily lost from birth onwards, a temporal loss of follicles that is different to that seen in other CO mutants. We find that this follicle loss in *Cntd1* mutants is dependent on the checkpoint kinase CHK2. Thus, in females, loss of *Cntd1* appears to result in phenotypes that are temporally disconnected from early and late CO mutants such as MutSγ and MutLγ, which show early prophase I disruption and ablation of ovary structure, and no prophase I disruption and an appearance of wildtype ovaries, respectively. These data suggest novel dual roles for CNTD1 in CO designation and faithful progression of oocytes into dictyate arrest at late pachynema, the latter being critical for establishing the ovarian reserve in female mice.

## INTRODUCTION

Meiosis is a specialized cellular division process that gives rise to haploid gametes for sexual reproduction. In contrast to mitosis, meiosis consists of two chromosome divisions preceeded by only a single round of DNA replication. During meiosis I, homologous parental chromosomes must find each other, pair and then segregate equally at the first meiotic division, resulting in up to two daughter cells that then enter meiosis II. The second division is reminiscent of mitosis in that it involves the segregation of sister chromatids that are already paired through the network of cohesion complexes that ensure their persistent connection. Thus, the faithful pairing and segregation of homologous chromosomes during meiosis I, followed by the accurate segregation of sister chromatids at meiosis II, are critical to ensuring the formation of euploid gametes. There appears to be distinct sexual dimorphism in the success of these events in mammalian meiosis, as underscored by the fact that that between 10-70% of human oocytes exhibit aneuploidy in comparison to only 3-5% of spermatocytes (Gruhn and Hoffmann, 2022; Gruhn et al., 2013; Hassold and Hunt, 2001; Hunt and Hassold, 2008; Morelli and Cohen, 2005; Nagaoka et al., 2012; Wang et al., 2017). Interestingly, but somewhat unsurprisingly, a majority of the nondisjunction events that lead to aneuploidy occur at the first meiotic division, arising from defects in crossover placement and designation, which are exacerbated by weakened cohesion between homologous chromosomes (Nagaoka et al. 2012; Hassold and Hunt 2001). This raises the question of how crossover regulation may differ between the sexes in mammals.

Segregation of homologous chromosomes at the first meiotic division is dependent on the unique events of prophase I: synapsis and recombination. The latter is initiated by the formation of hundreds of DNA double strand breaks (DSBs), each of which must be repaired with absolute fidelity prior to metaphase I entry. The majority of DSBs (∼90%) are repaired as non-crossovers (NCO) which do not involve exchange of flanking DNA sequence between homologous chromosomes. Remarkably, only a small remaining subset (10%) of the DSBs will be repaired as crossovers (CO) that are crucial for ensuring the persistence of homolog interactions until the first meiotic division. Given the importance of COs in ensuring persistent homolog interactions, therefore, it is not surprising that the distribution and frequency of COs across the genome must be tightly regulated to prevent abnormal chromosome segregation (Gray and Cohen 2016; Hunter 2015).

The majority of our knowledge of DSB repair and CO formation in mammalian meiosis comes from studies in male mice. In males, meiotic recombination involves a series of tightly regulated events downstream of DSB induction that begins with processing of DSBs to create single-stranded overhangs capable of invading an opposing homolog to test for homology. The initiation of DSBs is catalyzed by the topoisomerase-like protein SPO11 and its crucial partner proteins (Gray and Cohen, 2016; Keeney et al., 1997; Romanienko and Camerini Otero, 2000; Tran and Schimenti, 2019). Subsequently, the RecA homologs RAD51 and DMC1 facilitate the induction of homology search for either the repair of these DSBs between sister chromatids or homologous chromosomes (Cao et al. 1990; Cloud et al. 2012; Bishop et al. 1992; Hinch et al. 2020; Yoshida et al. 1998; MacQueen 2015). The latter are further processed into COs by a highly regulated network of pro-CO proteins.

The DNA mismatch repair MutS homologs, MSH4 (MutS Homolog 4) and MSH5 (MutS Homolog 5), which form a heterodimer known as MutSγ, associate with a subset of DSB repair intermediates in a process termed CO licensing (Edelmann et al. 1999; Kneitz et al. 2000; de Vries et al. 1999; Milano et al. 2019). Licensing also appears to require recruitment of RNF212 and RNF212B (Ring Finger protein 212 and 212B). (Condezo et al. 2024; Reynolds et al. 2013; Qiao et al. 2014; Qiao et al. 2018). Importantly, however, not all MutSγ/RNF212-loaded sites are destined to become COs in mammals. Instead, the final designation of COs requires the recruitment of a group of regulators including HEI10 (Human Enhancer of Invasion 10), CNTD1 (Cyclin N-terminal domain containing-1 protein), PRR19 (Proline-rich protein 19), and CDK2 (Cyclin-dependent kinase 2) (Bondarieva et al., 2020; Gray et al., 2020; Holloway et al., 2014; Palmer et al., 2020; Qiao et al., 2014; Reynolds et al., 2013; Ward et al., 2007), along with a heterodimer of the MutL homologs, MLH1 (MutL Homolog 1) and MLH3 (MutL Homolog 3), collectively named MutLγ. Together, these proteins mark the final subset of DSBs that will become COs of the major class I category (Edelmann et al., 1996; Kolas et al., 2005; Lipkin et al., 2002; Toledo et al., 2019; Woods et al., 1999). In mouse, class I COs represent 90-95% of all COs, and these accumulate across the genome at a frequency of 22-24 in males and 24-26 in females (Gray and Cohen, 2016; Lenzi et al., 2005). How a subset of the ∼150 MutSγ sites are designated to become COs is a major question in the field.

Previous studies in our lab revealed a critical role for CNTD1 in the process of crossover designation in male meiosis. CNTD1 was identified as an ortholog of the *C. elegans* Crossover Site-Associated-1 (COSA-1) (Yokoo et al., 2012). *cosa-1* mutant worms exhibit a high rate of inviable embryos and a significant increase in autosomal mis-segregation due to an absence of chiasmata, establishing *cosa-1* as a pro-CO factor (Yokoo et al., 2012). Interestingly, COSA-1 is a cyclin B-related protein, and recent studies have demonstrated that COSA-1 interacts physically with the cyclin-dependent kinase, CDK-2, to phosphorylate the C-terminus of MSH-5 to stabilize its association at sites of crossovers (Haversat et al. 2022; Zhang et al. 2021).

In mice, CNTD1 appears at pachynema of prophase I where it co-localizes with MutLγ and CDK2 (Gray et al., 2020; Holloway et al., 2014). Loss of CNTD1 in *Cntd1^-/-^* males results in the complete absence of epididymal spermatozoa leading to infertility (Gray et al., 2020; Holloway et al., 2014). During prophase I, spermatocytes from *Cntd1^-/-^*males show normal progression through leptonema and zygonema, including normal assembly of the synaptonemal complex and early DSB processing stages. By pachynema, however, spermatocytes from *Cntd1^-/-^* males fail to undergo normal CO designation, characterized by persistently elevated MutSγ focus frequency through to late pachynema, and failure to accumulate CO-associated MutLγ and CDK2 (Gray et al., 2020; Holloway et al., 2014). Interestingly, and unlike the situation for worm COSA-1, CNTD1 protein in the male mouse lacks a critical cyclin homology domain predicted to be encoded within the full-length genomic sequence, arising as an alternative translation start site within exon 3 (Gray et al., 2020). Accordingly, both yeast two-hybrid analysis and immunoprecipitation from testis extracts reveal that the testis CNTD1 is incapable of binding to CDK2 and thus does not act as a canonical cyclin during prophase I in the mouse, suggesting a different mechanism of action to that of COSA-1 (Gray et al., 2020; Haversat et al., 2022; Yokoo et al., 2012).

Given the sexual dimorphism in prophase I regulation observed in mammals, we were interested in investigating the role of CNTD1 in female meiosis. In contrast to the meiotic program in males, which initiates at sexual maturity and progresses continuously throughout life, meiosis in mammalian females begins during fetal development. More specifically, only prophase I is completed in fetal oocytes, then the oocytes pause in a unique stage called dictyate arrest just after diplotene, which is maintained until resumption of meiosis upon ovulation (Borum 1961). It is not until sexual maturity when meiosis I resumes only for a small cohort of oocytes. These select oocytes are arrested again at the end of metaphase II, and only at fertilization does the meiotic program complete (Morelli and Cohen, 2005). We find that *Cntd1* homozygous mutant females are sterile due to a failure to load MutLγ during pachynema of prophase I, resulting in an almost complete absence of chiasmata at metaphase I, similar to that seen for females lacking either MLH1 or MLH3 (Woods et al. 1999; Kan et al. 2008). Interestingly, however, the phenotypic consequence of this CO disruption differs in *Cntd1* mutant females from that in *Mlh1* or *Mlh3* homozygous mutant females: we observe a dramatic decrease in primordial follicles starting soon after birth in the absence of CNTD1, wherease loss of MLH1 or MLH3 results in normal primordial follicles through until adulthood (Edelmann et al. 1996; Lipkin et al. 2002). These subtle differences in outcomes for oocyte survival and subsequent folliculogenesis suggest that CNTD1 may play roles outside of the canonical CO designation pathway, most likely involving its cyclin-independent regulation of meiotic cell cycle events (Gray, 2020).

## RESULTS

### Oocytes from *Cntd1*^-/-^ female mice show a complete absence of class I crossovers

To determine the localization of CNTD1 in prophase I oocytes, we generated chromosome spreads from ovaries of mouse embryos at 18.5 days post-coitum (dpc) which corresponds to the developmental time point during which most of the germ cells are in pachynema of prophase I (Sun and Cohen 2013; Hwang et al. 2018; Borum 1961). We utilized two lines of mice, generated in our previous studies (Gray et al., 2020): the first bearing a null allele of *Cntd1*, referred to as *Cntd1^-/-^*, and the second harboring a tandem FLAG and HA-tagged variant of *Cntd1*, referred to as *Cntd1^HA/HA^*. The *Cntd1^HA^* allele behaves much like the wildtype allele, and thus, for all our experiments, we crossed both alleles to generate one experimental mouse strain, using *Cntd1^HA/HA^*mice as our wildtype controls, *Cntd1^HA/-^* mice as our heterozygous animals, and *Cntd1^-/-^*mice as our homozygous mutant test mice. Following chromosome spreading, oocyte nuclei were stained with antibodies against a component of the axial element, SYCP3 (Synaptonemal Complex Protein 3) and either the HA epitope for CNTD1 visualization (Fig. 1A-C) or MLH1 for class I CO maturation (Fig. 1E-G). We also performed co-localizaton of CNTD1 and MLH1 using primary antibodies for each that were raised in distinct host species (Suppl. Fig. 1). CNTD1 was detected as discrete foci localized to the synaptonemal complex in pachytene-staged oocytes in a pattern and frequency reminiscent of the terminal crossover factors, MLH1 and MLH3 (Fig. 1A,B,D). CNTD1^HA^ was observed on spreads from both *Cntd1*^HA/HA^ and *Cntd1*^HA/-^ (n=157 nulei; mean ± S.E.M; 27.34 ± 0.3097 and n=66 nuclei; 26.06 ± 0.4083, respectively). CNTD1^HA^ foci were significantly decreased in *Cntd1*^HA/-^ compared to *Cntd1*^HA/HA^ (p=0.0185, Mann-Whitney test) (Fig. 1D). As expected, *Cntd1*^-/-^ oocytes (n=44 nuclei) did not show the presence of CNTD1^HA^ on chromosome spreads, resulting in a significant decline compared to *Cntd1^HA/-^*and *Cntd1^HA/HA^* oocytes (p<0.0001 and p<0.0001, respectively; Mann-Whitney test) (Fig. 1C). This result is consistent with our published work characterizing the function of CNTD1 in mammalian spermatogenesis (Gray et al., 2020).

**Figure 1.**
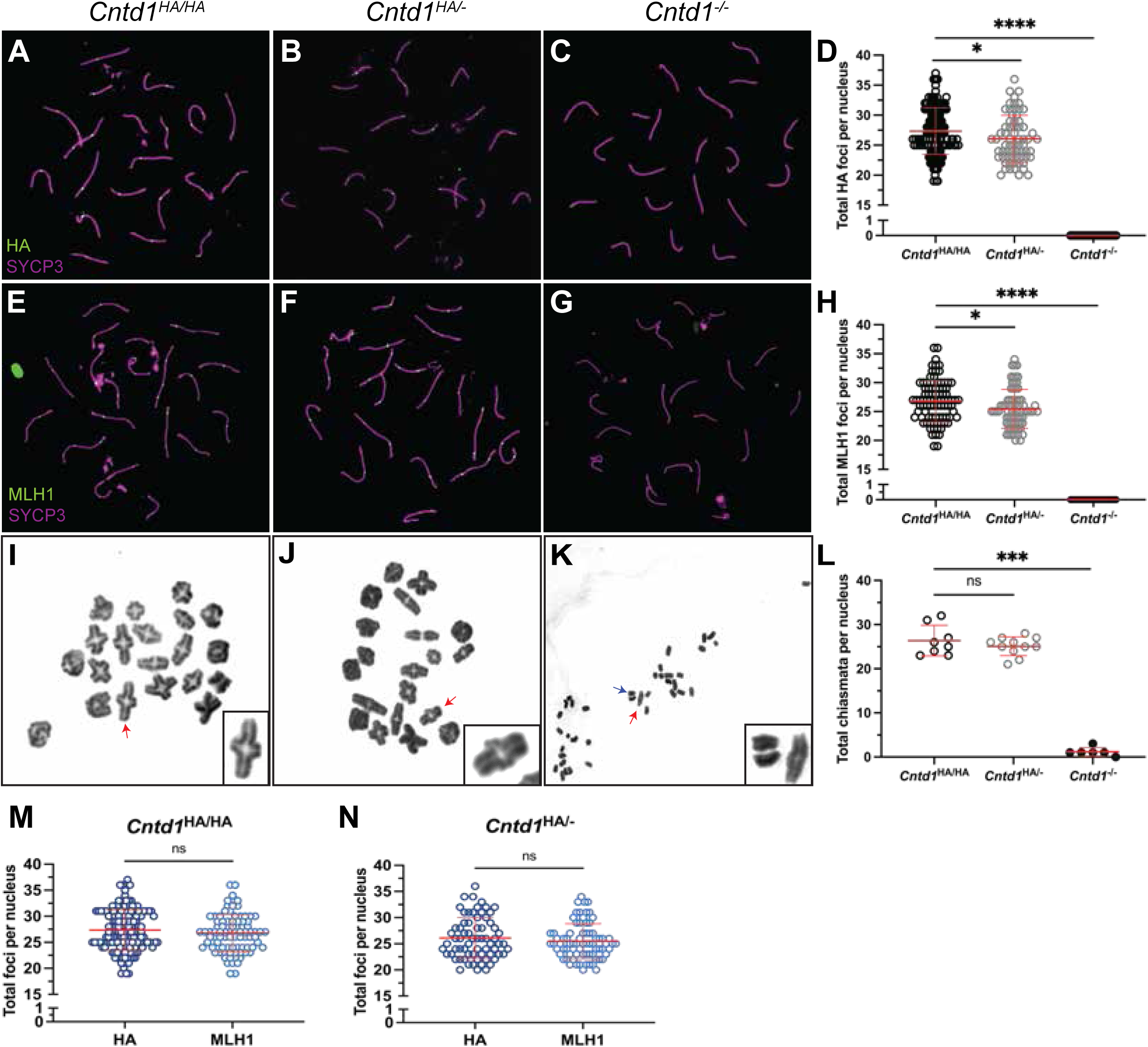
*Cntd1*^-/-^ females lack class I crossovers and show a significant increase in univalent chromosomes. Localization of CNTD1-HA (green) and SYCP3 (magenta) in pachytene oocytes from (A) *Cntd1*^HA/HA^, (B) *Cntd1*^HA/-^, and (C) *Cntd1*^-/-^ fetuses using antibodies against each protein raised in rabbit and mouse, respectively. Localization of MLH1 (green) and SYCP3 (magenta) in pachytene oocytes from (E) *Cntd1*^HA/HA^, (F) *Cntd1*^HA/-^, and (G) *Cntd1*^-/-^ females using antibodies against each protein raised in mouse and rabbit, respectively. (D) Quantification (average foci ± SEM) of HA foci in oocytes from *Cntd1*^HA/HA^ (n=157), *Cntd1*^HA/-^ (n=66), and *Cntd1*^-/-^ (n=47 nuclei) females and (H) MLH1 foci in oocytes from *Cntd1*^HA/HA^ (n=87), *Cntd1*^HA/-^ (n=73) and *Cntd1*^-/-^ (n=47 nuclei) females. Diakinesis preps with quantification in graphs, arrows indicate bivalents (red) and univalent (blue). (I) *Cntd1*^HA/HA^, (J) *Cntd1*^HA/-^, and (K) *Cntd1*^-/-^ chiasmata visualized with Giemsa staining. (L) Quantification (average chiasmata ± SEM) of chiasmata for each genotype (*Cntd1*^HA/HA^ n=8 cells, *Cntd1*^HA/-^ n=11 cells, and *Cntd1*^-/-^ n=9 cells). (M and N) Statistical comparison of HA and MLH1 foci in *Cntd1*^HA/HA^ and *Cntd1*^HA/-^ oocytes. Mean and standard error mean lines are in red. All experiments utilized n ≥ 3 ovary pairs per genotype. A Mann-Whitney test was utilized to test for statistical significance between genotypes. P-values are as follows, p<0.05 (*), p<0.001 (**), p<0.0002 (***), p<0.0001 (****).

To explore the distribution of class I crossover events across the genome in the absence of CNTD1, chromosome spreads were prepared from pups of different genotypes, and stained with antibodies against SYCP3 and MLH1. MLH1 foci were observed in both *Cntd1^HA/HA^* and *Cntd1^HA/-^*oocytes, but not *Cntd1^-/-^* oocytes (Fig. 1E, F, H; n=87 nuclei; mean ± S.E.M; 26.76 ± 0.3841, n=73 nuclei; 25.47 ± 0.3941, n=47 nuclei, respectively). *Cntd1^HA/-^*oocytes also exhibited a significant decrease in the number of MLH1 foci compared to *Cntd1^HA/HA^*oocytes (p=0.0112). There is no difference in the quantity of CNTD1^HA^ foci compared to MLH1 foci in both *Cntd1^HA/HA^* and *Cntd1^HA/-^* oocytes (Fig. 1M, N; p=0.2848 and p=0.4310, respectively).

The significant decline in both CNTD1^HA^ and MLH1 foci in *Cntd1^HA/-^* oocytes could be due to the dosage of *Cntd1*, which has been observed in other pro-crossover factors such as *Hei10* and *Rnf212* (Qiao et al. 2014; Reynolds et al. 2013). Similar to our findings in *Cntd1*^-/-^ males, the absence of MLH1 indicates that CNTD1 is either acting upstream of MutLγ, or that CNTD1 is necessary for MutLγ accumulation at sites of COs.

To investigate co-localization of CNTD1 and MutLγ, we used antibodies against HA in conjunction with our custom-made antibodies against MLH3 (Gray et al., 2020). On average, ∼89% and ∼93% of CNTD1^HA^ foci were found to co-localize with MLH3 at pachynema in spreads from Cntd1*^HA^*^/*HA*^ and *Cntd1*^HA/-^ oocytes, respectively. Conversely, ∼91% and ∼87% of MLH3 foci localized with CNTD1^HA^ in *Cntd1*^HA/HA^ and *Cntd1*^HA/-^ oocytes, respectively (Suppl. Fig 1). The close co-localization pattern of CNTD1 with MLH3 lends evidence to the function of the former as a class I crossover regulator. This result is also supported by the similar number of MLH1 and CNTD1 foci at pachynema (Fig 1D, H).

### Crossover maturation is disrupted in *Cntd1*^-/-^ females

At the end of prophase I, COs manifest into physical linkages holding homologous chromosomes together, known as chiasmata. We found that chiasmata formation was normal in *Cntd1*^HA/HA^ and *Cntd1*^HA/-^ oocytes (red arrows, Fig 1I, J, L; n=8, 26.38 ± 1.224 and n=11 cells, 25.09 ± 0.639, respectively). However, *Cntd1*^-/-^ oocytes showed a significant decline in chiasmata formation compared to *Cntd1*^HA/HA^ and *Cntd1*^HA/-^ oocytes resulting in an increase of univalent chromosomes (blue arrow, Fig. 1K; n=6 cells, p=0.0007 and p<0.0001, respectively). However, *Cntd1*^-/-^ oocytes retained on average 1.167 ± 0.4014 chiasmata per cell (red arrow, Fig. 1L, K), reflecting an approximately 95% loss of COs, presumably indicative of a functional class II crossover pathway, albeit at a slightly reduce rate of use to that seen in male meiosis (Holloway et al., 2014, 2008). Taken together, these results indicate that *Cntd1* is essential for class I crossover formation, which is consistent with the model that CNTD1 is involved in regulating MLH1 placement, but not essential for class II crossover formation as evidenced by a persistence of a small number of residual chiasmata.

### CNTD1 is expressed in a truncated isoform lacking critical cyclin homology domains

Like its nematode ortholog, COSA-1, the predicted “full-length” isoform of CNTD1 is predicted to act as a cyclin due to the presence of three cyclin homology regions at amino acid positions 57-88, 117-135, and 140-180 (Gray et al., 2020) (Fig. 2B). However, *in silico* analysis of the *Cntd1* coding sequence shows a highly conserved second methionine at amino acid position 86 (in mouse), which may result in an alternative translation start site to produce a shorter form of CNTD1 that is 249 amino acids in length (compared to the 334 amino acid length of the predicted full-length protein). The smaller CNTD1 protein was confirmed *in* vivo with Western blots in multiple studies (Bondarieva et al. 2020; Gray et al. 2020). CNTD1 lacks the ability to interact with crossover associated CDK2, its putative kinase interactor, in both yeast two-hybrid assays (when the shorter CNTD1 sequenced is used) and co-immunoprecipitations from testis lysate (Bondarieva et al., 2020; Gray et al., 2020). Importantly, smaller CNTD1 is the only form of CNTD1 found in mouse spermatocytes, or indeed in any tissue in male mice (Gray et al., 2020) or in germ cells from men, dogs and cats (Wolff and Cohen, unpublished observations). Therefore, we tested whether CNTD1 exists as the same short form in mouse oocytes, or perhaps is encoded by an alternative isoform of CNTD1 in female mice, possibly providing an explanation for the observed sexual dimorphism in prophase I outcomes between males and females.

**Figure 2.**
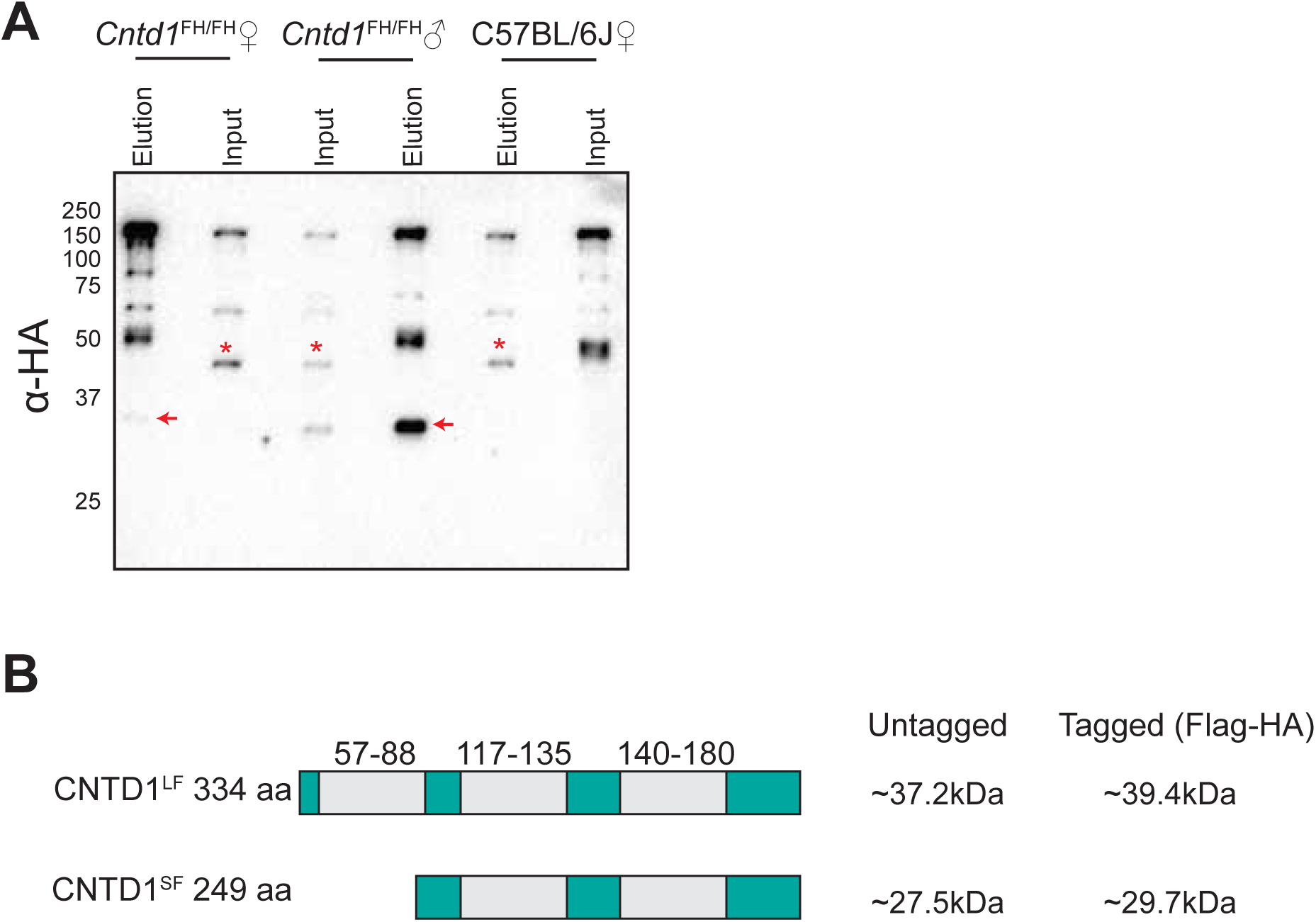
CNTD1 exists in a short-form in fetal oocytes. Co-immunoprecipitation with anti-HA antibody in fetal ovary and adult testis lysate. (A) Western Blot against HA from IPs performed in 18.5dpc whole ovary lysate from *Cntd1*^HA/HA^ and C57BL/6J females, and whole testis lysate from *Cntd1^HA^*^/*HA*^ adult males. Approximately 40-100 ovaries from n ≥ 20 fetuses were utilized per sample. Red arrows indicate the predicted mass of CNTD1 (∼29.7kDa) in the elutions from fetal ovaries and adult testes from *Cntd1*^HA/HA^ animals. Red astriks indicate non-specific bands detected by the HA antibody. (B) *In silico* prediction of the long form and short form of CNTD1 with cyclin homology domains in light grey and predicted molecular weight with and without Flag-HA epitope tags to the left of the diagram.

We performed a co-immunoprecipitation (co-IP) with antibodies against HA to isolate CNTD1-HA from fetal ovary cell lysates collected at 18.5 dpc and then probed via Western blot (WB) with anti-HA antibody to determine which isoform of CNTD1 was expressed in oocytes. CNTD1-HA was present in the IP elutions at the same predicted molecular weight (∼29.7 kDa) in oocytes (*Cntd1^HA/HA^* ♀) as in spermatocytes (*Cntd1^HA/HA^*♂) (Fig. 2A, red arrows). As expected, we did not find HA-tagged CNTD1 in fetal whole ovary lysate from C57BL/6J females (C57BL/6J ♀). A non-specific band at ∼40kD is present in both the HA samples and in the C57BL/6J control (red asterisks), therefore this is not CNTD1-HA. Taken together, these data support our previous study showing CNTD1 exists *in vivo* in an isoform lacking the first cyclin homology domain, and suggests that CNTD1 is functioning via a cyclin independent mechanism in mammalian oocytes.

### Absence of *Cntd1* results in sterility and loss of primordial follicles

In males, disruption of factors essential for class I CO formation and distribution results in complete sterility and manifests in a phenotype characterized by gross morphological defects in gonadal tissues, including a significant decrease in testis size coupled with a significant loss of meiocytes and loss of post meiotic cells in testis tissue sections (Bondarieva et al., 2020; Gray et al., 2020; Holloway et al., 2014). Morphological effects appear to be bifurcated in females where there are two distinct phenotypes. The first is observed in mutants that affect early DSB repair events, and leads to a significant reduction in ovarian size and lack of folliculogenesis resulting from disruption of meiotic prophase prior to pachynema. This phenotypic category is represented by mouse mutants lacking genes such as *Dmc1, Msh4, and Msh5* (Edelmann et al., 1999; Kneitz et al., 2000; Pittman et al., 1998; Yoshida et al., 1998). The second phenotypic category is characterized by later prophase I disruption from pachynema onwards, resulting in normal ovarian morphology and follicle composition, including genes such as *Mlh1*, *Hei1*0, and *Prr19* (Edelmann et al. 1996; Ward et al. 2007; Bondarieva et al. 2020). Due to the accumulation of CNTD1 in pachynema, and its co-localization with other pro-crossover factors we hypothesized a similar temporal phenotype for *Cntd1* females to those lacking *Mlh1, Hei10,* and *Prr19*, namely no loss of follicle structures or ovarian morphology in adult mice, but producing oocytes that have few to no crossovers.

To explore the consequences of CNTD1 dysfunction in the oocyte for ovarian development and folliculogenesis, we investigated fertility, ovarian morphology, and follicle populations using *Cntd1*^HA/HA^ animals as our wildtype control cohort together with *Cntd1*^HA*/-*^ and *Cntd1^-/-^* females. *Cntd1*^-/-^ females were unable to produce litters after approximately three months housed with a *Cntd1*^HA/HA^ male (Suppl. Fig. 2), confirming that these animals are infertile. *Cntd1*^HA*/-*^ females produced on average 1 fewer offspring compared to *Cntd1*^HA/HA^ females (Suppl Fig. 2). To determine if ovaries from *Cntd1*^-/-^ females exhibit early or late prophase I disruption, we quantitated the ovarian follicular composition from pre-pubertal post-natal day 1 (PND 1), post-natal day 14 (PND 14), post-natal day 28 (PND 28), and adult (8-12 week old) females (n≥3 for each time point), documenting the follicular stages from primordial (early) follicles through antral (late pre-ovulatory) follicles. Primordial follicles contain the dictyate arrested prophase I oocyte surrounded by one layer of granulosa cells and represent the finite pool of germ cells and reproductive potential of the individual. Maturation to primary through antral follicles are only induced upon sexual maturity and these follicle types are characterized by layers of granulosa and theca cells, which secrete hormones which facilitate this maturation process (Epifano and Dean 2002). Prematurely exhausted primordial follicle pools can be indicative of defects in oogenesis prior to the formation of primordial follicles.

At PND1 the combined number of germ cell cysts and primordial follicles in *Cntd1*^HA/HA^, *Cntd1*^HA/-^, and *Cntd1*^-/-^ were not significantly changed between *Cntd1*^HA/HA^ and *Cntd1*^HA/-^, and *Cntd1*^HA/HA^ and *Cntd1*^-/-^ (Fig. 3A-D; mean number of follicles ± SEM 460 ± 56.20, 423.3 ± 61.94, and 238.3 ± 78.01, respectively, p=0.6839 and p=0.0891, respectively). The first stage of follicular growth is represented by the appearance of primordial staged follicles, which are apparent at the PND14 timepoint onwards, and these exhibited a significant decrease in *Cntd1*^-/-^ females compared to *Cntd1*^HA/HA^ littermate controls (258.33 ± 36.55 and 1595.0 ± 172.0 follicles per ovary, respectively; p<0.01, Fig. 3E-H). There was no difference in the number of primordial follicles between *Cntd1*^HA/HA^ and *Cntd1*^HA/-^ animals (Fig. 3H; 1595.0 ± 172.0 and 1249 ± 191.7, p=0.2928). The number of secondary and antral-staged follicles were unchanged in *Cntd1*^HA/HA^, *Cntd1*^HA/-^, and *Cntd1*^-/-^ ovaries (Fig. 3H).

**Figure 3.**
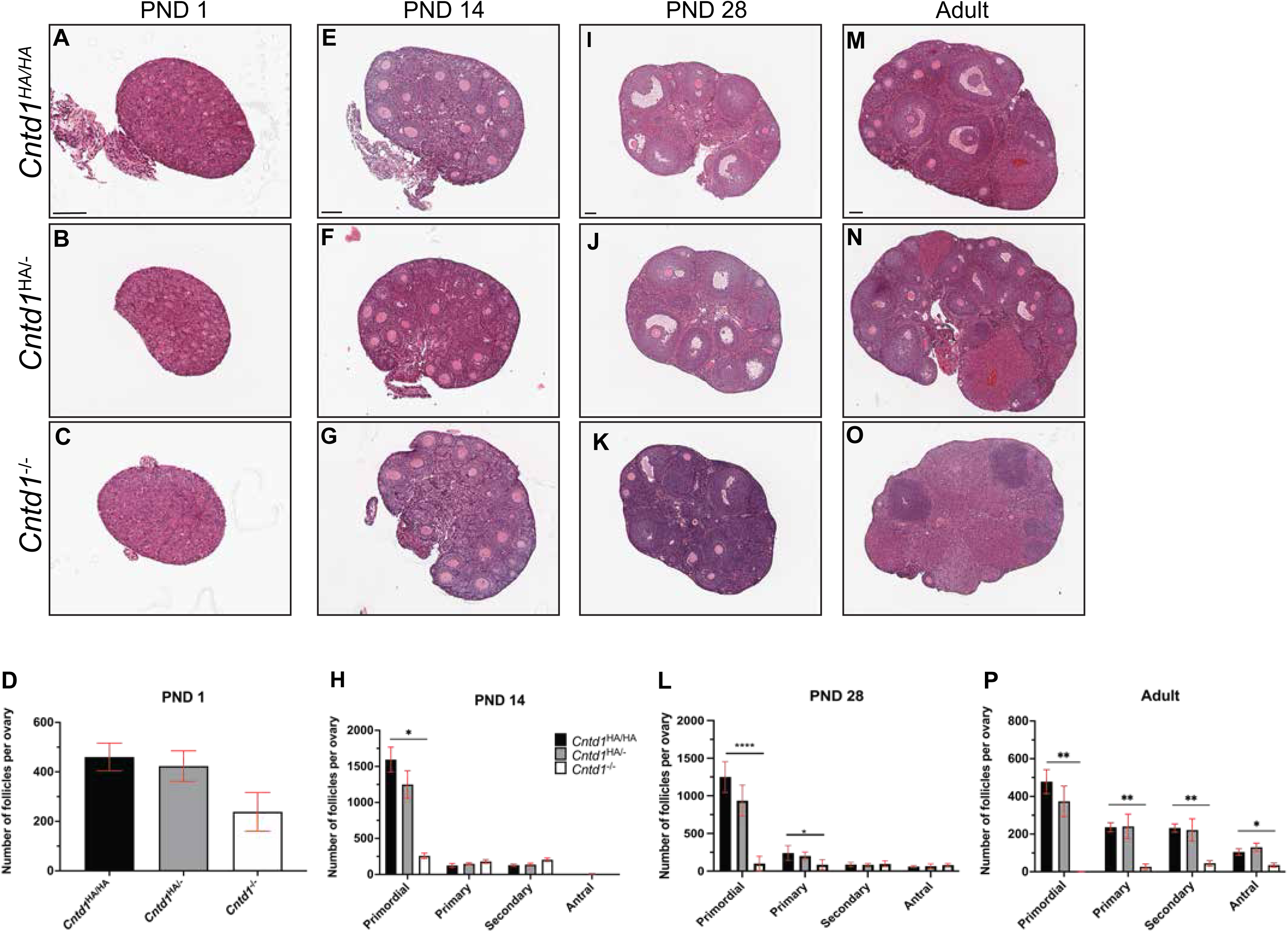
Ovarian histology in *Cntd1*^HA/HA^, *Cntd1*^HA/-^, and *Cntd1*^-/-^ females at different postnatal ages reveals a significant increase in the loss of primordial follicles. **(A-O)** Histological sections of ovaries from *Cntd1*^HA/HA^, *Cntd1*^HA/-^, and *Cntd1*^-/-^ female mice at post-natal day (PND) 1 (A-C), 14 (E-G), 28 (I-K), and adult (8-10 week) (M-O). Paraffin-embedded tissues were sectioned at 5 μm and stained with Hematoxylin and Eosin. Scale bars are equal to 100μm. Quantification (average follicle number ± SEM) of primordial, primary, secondary, and antral follicles in ovaries at PND1 (D), PND14 (H), PND28 (L), and adult (P) ages from *Cntd1*^HA/HA^, *Cntd1*^HA/-^, and *Cntd1*^-/-^ female mice. For each time point, follicles from n≥3 ovaries were quantified. Mean and standard error mean lines are in red. A Mann-Whitney test was utilized to test for statistical significance. P-values are as follows, p<0.05 (*), p<0.001 (**), p<0.0002 (***), p<0.0001 (****).

At PND28 we observed no change in the average number of primordial follicles found in *Cntd1*^HA/HA^ compared to *Cntd1*^HA/-^ ovaries, but there was a change in *Cntd1*^-/-^ compared to *Cntd1*^HA/HA^ ovaries (Fig. 3I-L; mean primordial follicles per ovary ± SEM; 1250 ± 103.2, 935.7± 78.6 86 ± 33.32, and 100.0 ± 49.03, respectively; p=0.0727 and p=0.0286, respectively). The average number of primary follicles was not changed in ovaries from *Cntd1*^HA/HA^ animals compared to *Cntd1*^-/-^ (Fig. 3L; 241.3 ± 48.96 and 86.25 ± 33.32 follicles per ovary, respectively; p=0.0571). There was no difference in the number of primary follicles between *Cntd1*^HA/HA^ and *Cntd1*^HA/-^ ovaries (Fig. 3L; 241.3 ± 48.96 and 200 ± 20.64 follicles per ovary; p=0.6485).

In adult *Cntd1*^HA/HA^ ovaries, we observed the following distribution of follicle types (mean ± SEM): 478.33 ± 63.96 primordial, 236.66 ± 24.28 primary, 232.5 ± 22.13 secondary, and 105 ± 17.61 antral follicles per ovary (Fig. 3M, P). *Cntd1*^HA/-^ ovaries had: 374.2 ± 81.79 primordial, 240.8 ± 65.39 primary, 221.7 ± 59.3 secondary, and 130 ± 21.64 antral follicles per ovary (Fig. 3N, P). By contrast, *Cntd1*^-/-^ ovaries contained the following ratios of follicle types: 1.250 ± 1.250 primordial, 26.25 ± 15.05 primary, 45.00 ± 14.86 secondary, and 35 ± 11.37 antral follicles per ovary, with significant declines in all follicular stages relative to *Cntd1*^HA/HA^ ovaries (Fig. 3O, P; p<0.05 for all follicle types). There was no significant difference between total follicle numbers in *Cntd1*^HA/HA^ and *Cntd1*^HA/-^ ovaries for any of the follicle types (Fig. 3P).

Taken together, these data suggest that the few primordial follicles that are able to progress through folliculogenesis after birth can then undergo normal development, however the oocytes are inviable. The loss of primordial follicles in the absence of CNTD1 only starts to impact primary, secondary and antral follicles from PND 28 onwards. These data indicate that oocytes from *Cntd1*^-/-^ mice start to undergo apoptosis from birth and before follicle formation occurs, at some point between the end of prophase I and before PND 14, and that this continues in a dramatic fashion as the mice progress towards adulthood. Given the temporal accumulation of CNTD1 and the phenotype of *Cntd1* deletion is similar to late CO designation mouse mutants, such as *Mlh1, Hei10,* and *Prr19*, it was surprising to observe early primordial follicle loss in *Cntd1* mutants, which is temporally distinct to these other CO designation mutants.

### *Cntd1*^-/-^ oocytes exhibit elevated unrepaired DSBs

Due to the premature loss of primordial follicles in *Cntd1*^-/-^ ovaries, we were interested in investigating the mechanism by which these oocytes were being lost. Incomplete synapsis of homologous chromosomes and unrepaired DSBs during meiotic prophase I in oocytes trigger the synapsis and DSB checkpoints (e.g., Dual-Checkpoint Model), respectively, leading to oocyte apoptosis in the absence of sufficient repair (Bolcun Filas et al., 2014; Ravindranathan et al., 2022; Rinaldi et al., 2020, 2017). To assess if early DSB repair and recombination events are affected in *Cntd1^-/-^* oocytes, we used an antibodies against the RecA homolog, RAD51, and γH2A.X to stain spread chromosomes in pachytene-staged oocytes in all *Cntd1* genotypes (Fig. 4A-L). All genotypes showed canonical staining patterns in which RAD51 accumulated as expected in zygotene-staged oocytes (Fig. 4D-F) and was almost completely absent by pachynema (Fig. 4G-I), indicating normal progression of DSB repair past the strand invasion stages. Upon analysis of RAD51 foci at pachynema, however, *Cntd1^HA^*^/*HA*^ showed a modest but significant increase in RAD51 focus counts in comparison to *Cntd1*^-/-^ oocytes (Fig. 4M; mean foci ± S.E.M; n=83 oocytes, 3.506 ± 1.184 and n=44 oocytes 4.659 ± 1.663, respectively; p=0.0173). There was no change in the number of foci between *Cntd1^HA^*^/*HA*^ and *Cntd1^HsA/-^* (Fig. 4M; 2.891 ± 1.068, respectively; p=0.7701).

**Figure 4.**
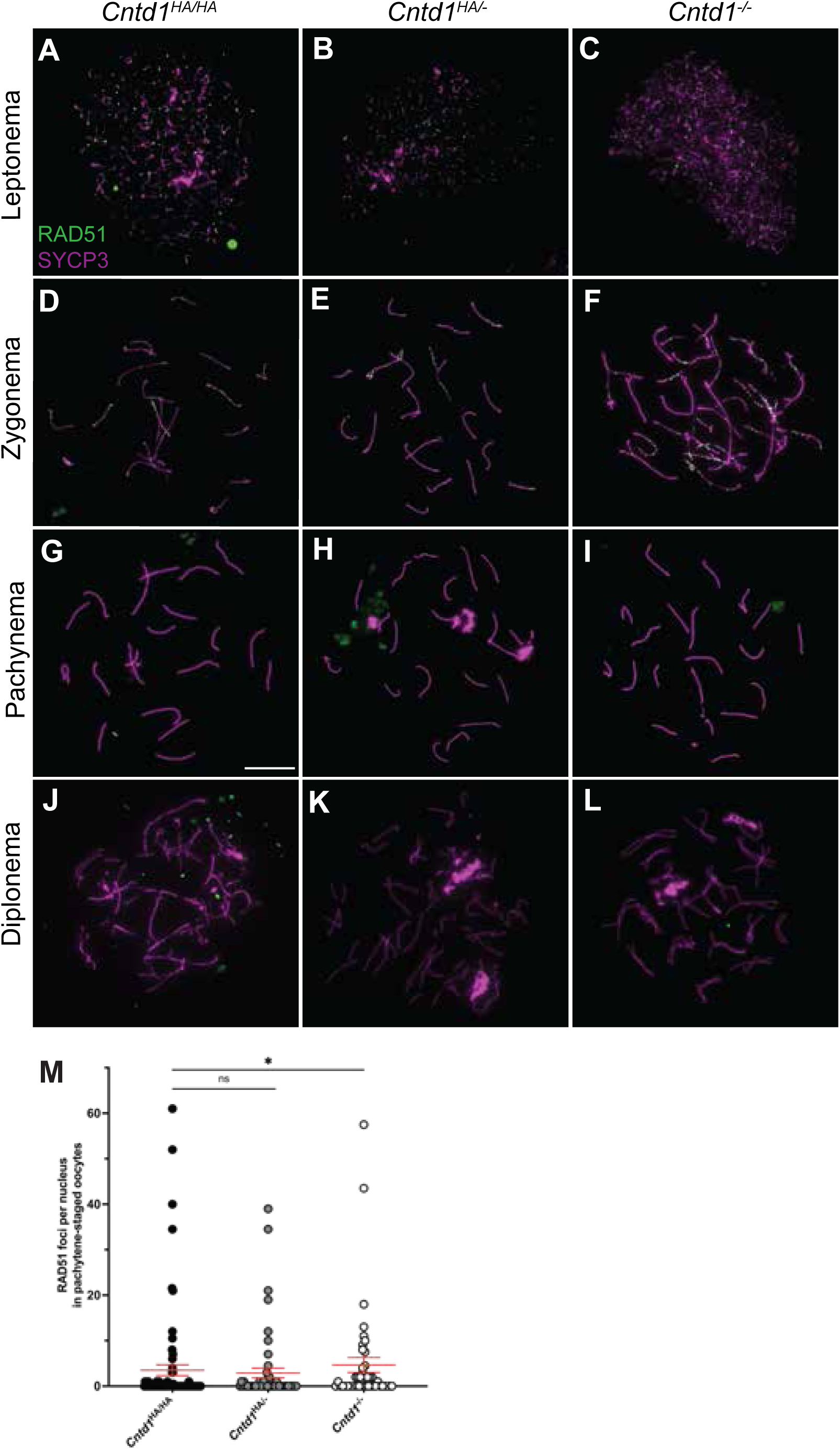
Early DSB repair in *Cntd1*^-/-^ oocytes is normal. Localization of RAD51 (green) and SYCP3 (magenta) on chromosomes spreads from *Cntd1*^HA/HA^, *Cntd1*^HA/-^, and *Cntd1*^-/-^ oocytes at leptonema (A-C), zygonema (D-F), pachynema (G-I), and diplonema (J-L). Using antibodies against each protein raised in rabbit and mouse, respectively. (M) Quantification of RAD51 in pachytene-staged cells, *Cntd1*^HA/HA^ (n=78 nuclei), *Cntd1*^HA/-^ (n=51 nuclei), *Cntd1*^-/-^ (n=44 nuclei). n ≥3 mice were used and data was analyzed using Mean and standard error mean lines are in red. A Mann-Whitney test was utilized to test for statistical significance. P-values are as follows, p<0.05 (*), p<0.001 (**), p<0.0002 (***), p<0.0001 (****).

We next investigated γH2A.X, which serves as another marker of unrepaired breaks and in its unphosphorylated state as H2A.X, is a target of surveillance kinases such as ATR and ATM in the DNA damage response pathway (Collins et al. 2020). We measured the nuclear intensity of γH2A.X signal in pachytene oocytes from *Cntd1*^-/-^ and *Cntd1*^HA/HA^ embryos and observed that the former was markedly reduced compared to the latter (Fig. 5A, C, J; n=40 oocytes, n=60 oocytes, respectively; p=0.0003). γH2A.X was also reduced in *Cntd1*^HA/-^ oocytes compared to *Cntd1*^HA/HA^ littermate controls (Fig. 5A, B, J; n=149 oocytes, p=0.0315).

**Figure 5.**
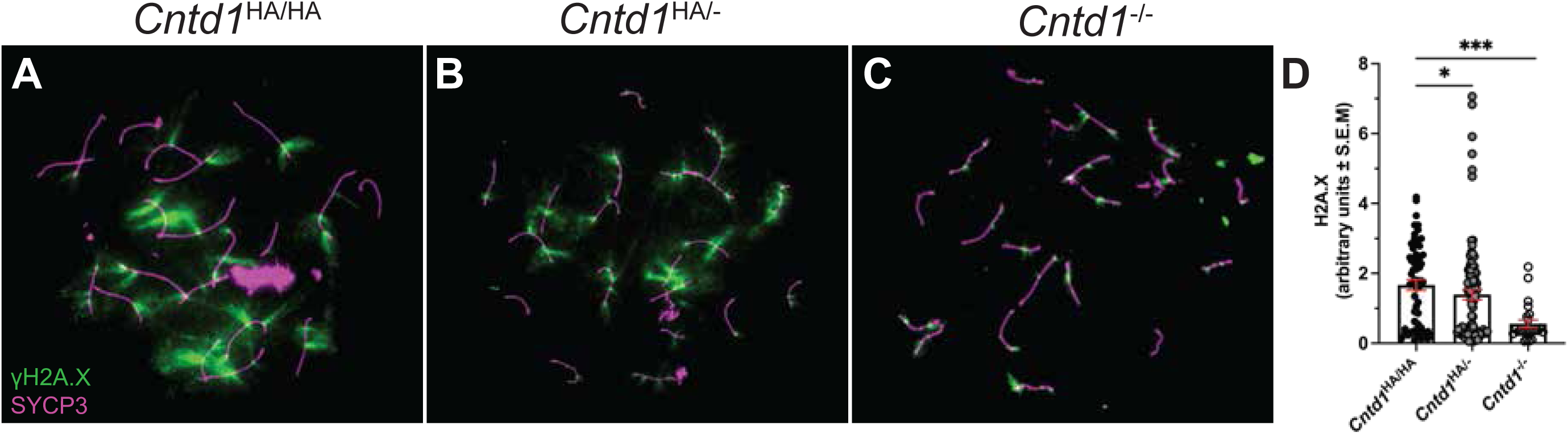
γH2A.X intensity is reduced in *Cntd1*^-/-^ oocytes. Localization of γH2A.X (A-C) with SYCP3 (magenta) on chromosomes spreads from *Cntd1*^HA/HA^, *Cntd1*^HA/-^, and *Cntd1*^-/-^ oocytes at pachynema. (D) Quantification of the intensity of γH2A.X normalized to DAPI in pachytene-staged cells, *Cntd1*^HA/HA^ (γH2A.X n=60 nuclei), *Cntd1*^HA/-^ (γH2A.X n=94 nuclei), and *Cntd1*^-/-^ (γH2A.X n=24 nuclei). n ≥3 fetuses were used and data was analyzed using Mean and standard error mean lines are in red. A Mann-Whitney test was utilized to test for statistical significance. P-values are as follows, p<0.05 (*), p<0.001 (**), p<0.0002 (***), p<0.0001 (****).

Although there was a mild increase in the number of RAD51 foci in pachynema-staged oocytes, this increase is most likely not sufficient to trigger the DSB portion of the dual checkpoint due to previous reports indicating that oocytes can withstand ∼10 unrepaired DSBs before induction of the checkpoint (Ravindranathan et al. 2022). The decrease in γH2A.X intensity in *Cntd1* mutant pachytene-staged oocytes could indicate either a deficiency in repairing DSBs or acceleration in repairing DSBs. We speculate that aberrant/failed DSB repair could contribute to the loss of oocytes lacking CNTD1 in prophase I.

### *Cntd1*^-/-^ oocytes have a premature acceleration through prophase I

Previous work from our lab showed that spermatocytes from *Cntd1*^-/-^ males show no overt defects in SC formation and synapsis (Gray et al., 2020; Holloway et al., 2014). To investigate whether such defects in SC assembly could explain the early loss of primordial follicles in *Cntd1*^-/-^ females, we analyzed synapsis in prophase I oocytes from 18.5dpc *Cntd1*^-/-^ females using antibodies against the synaptonemal complex markers, SYCP1 (Synaptonemal complex protein-1) and SYCP3. Beginning in zygonema, SYCP1 transverse filaments zip the central and axial elements of the synaptonemal complex together (de Vries et al., 2005). Full synapsis is the defining feature of pachynema in all meiotic organisms. Defects in the ability for these proteins to assemble properly results in quality control checkpoint activation and subsequent apoptosis at the end of prophase I (Hamer et al., 2008; Yuan et al., 2000).

We observed normal SYCP1 localization in leptotene through diplotene-staged oocytes from *Cntd1*^HA/HA^, *Cntd1*^HA/-^, and *Cntd1*^-/-^ pups at 18.5 dpc (Fig. 6A-L). Although all substages of prophase I were observed in *Cntd1*^-/-^ oocytes, the distribution of these substages were altered compared to *Cntd1*^HA/HA^ oocytes. To interrogate prophase I progression, we quantified the first 100 oocytes per slide for each genotype from 18.5dpc fetal ovaries and sub-staged oocytes based on the staining pattern of SYCP1 and SYCP3 (see Materials and Methods). Pachytene-staged oocytes in *Cntd1*^HA/HA^ animals were present at ∼70%, but only observed at ∼13% in *Cntd1*^-/-^ (Fig. 6M). This alteration in meiotic progression continued, as ∼20% of *Cntd1*^HA/HA^ oocytes were at diplotene in contrast to ∼79% of *Cntd1*^-/-^ oocytes (Fig. 6M).

**Figure 6.**
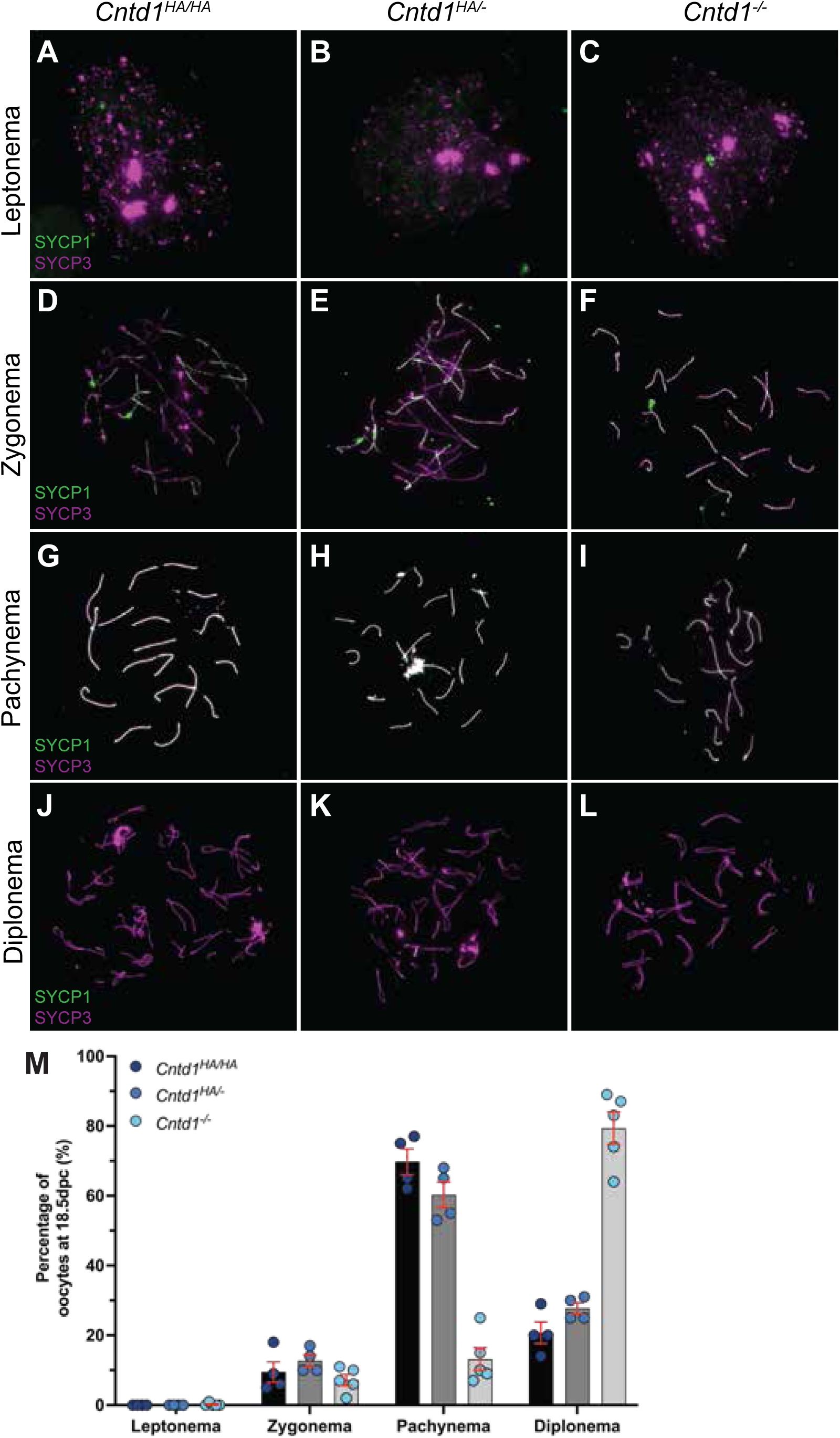
Progression through prophase I in *Cntd1*^-/-^ oocytes is altered. Localization of SYCP1 (green) and SYCP3 (magenta) on chromosomes spreads from *Cntd1*^HA/HA^, *Cntd1*^HA/-^, and *Cntd1*^-/-^ oocytes at leptonema (A-C), zygonema (D-F), pachynema (G-I), and diplonema (J-L). Using antibodies against each protein raised in rabbit and mouse, respectively. (M) Analysis of prophase I sub-stages in 18.5dpc *Cntd1*^HA/HA^ (black bar, dark blue circles), *Cntd1*^HA/-^ (dark grey, medium blue circles), and *Cntd1*^-/-^ (light grey, light blue circles) oocytes. n≥3 biological replicates per genotype, indicated by colored circles.

We next investigated the localization of SKP1, which is a component of the SCF (SKP1-Cullin-F-box complex) and is essential for cell cycle progression during mouse meiosis (Guan et al. 2020; Guan et al. 2022). Our previous studies in mouse spermatocytes found that CNTD1 interacts with components of the SCF complex and *Cntd1* mutant spermatocytes show a prophase I to metaphase I transition defect (Gray et al. 2020), similar to *Skp1cKO* mutants (Guan et al. 2020). More recent characterization of SKP1 during meiosis has shown it to be essential for synapsis in mammalian oocytes and spermatocytes (Guan et al. 2022). In *Cntd1*^HA/HA^ oocytes, we observed normal SKP1 localization to SCs, however, this localization appeared to be completely lost in *Cntd1*^-/-^ oocytes (Fig. 7A-C). We next measured the intensity of SKP1 in *Cntd1* oocytes and observed that compared to *Cntd1*^HA/HA^, *Cntd1*^-/-^ but not *Cntd1* ^HA/-^ showed a significant reduction in the intensity of SKP1 (Fig. 7D; p<0.0001 and p=0.5569, respectively).

**Figure 7.**
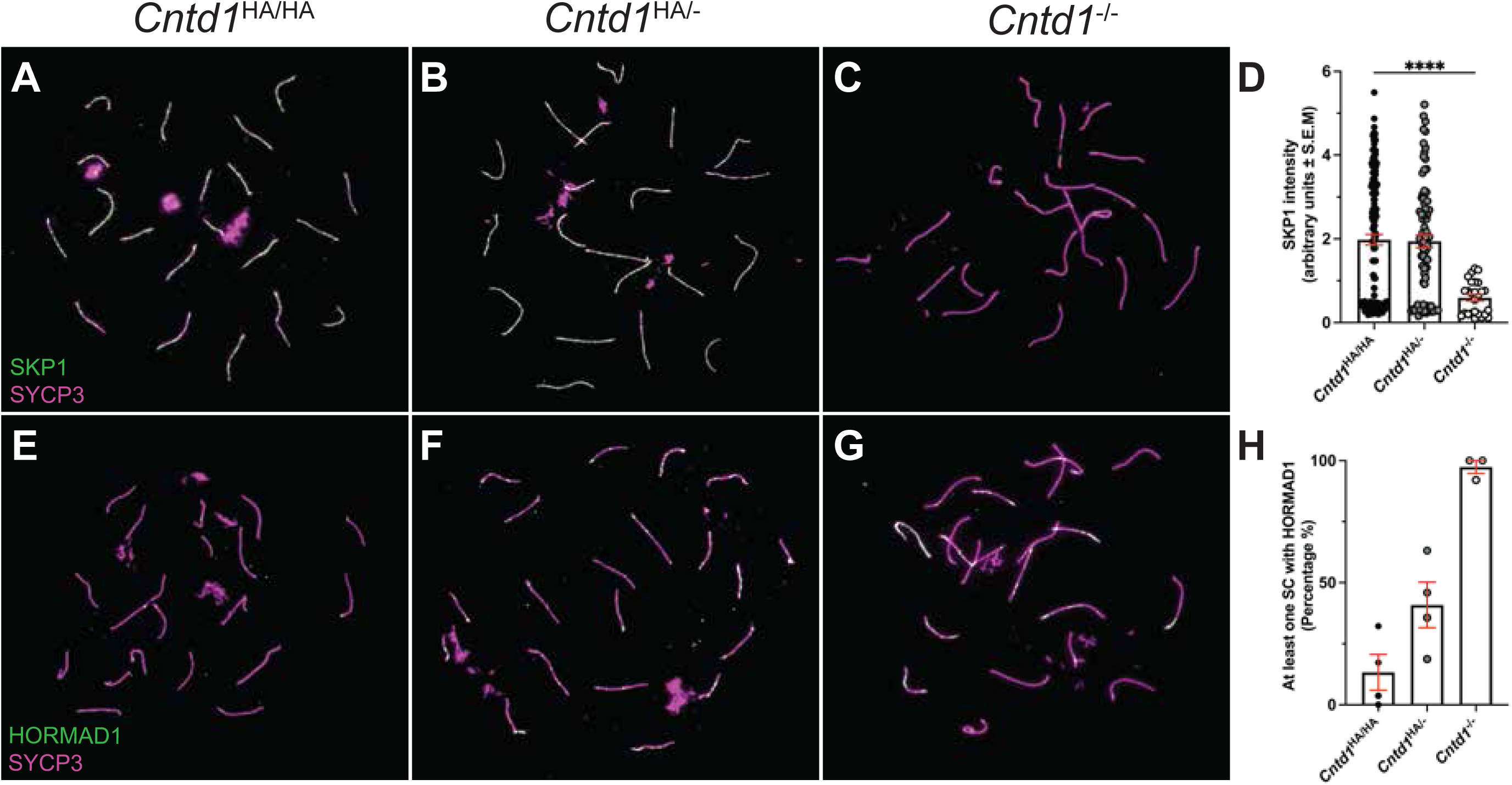
HORMAD1 and SKP1 are abberant in *Cntd1*^-/-^ oocytes. Localization of SKP1 (A-C) and HORMAD1 in green (E-G) with SYCP3 (magenta) on chromosomes spreads from *Cntd1*^HA/HA^, *Cntd1*^HA/-^, and *Cntd1*^-/-^ oocytes at pachynema. (D and H) Quantification of the intensity of each antibody of interest normalized to DAPI in pachytene-staged cells, *Cntd1*^HA/HA^ SKP1 n=142 nuclei and HORMAD1 n=132 nuclei), *Cntd1*^HA/-^ (SKP1 n=81 nuclei and HORMAD1 n=149 nuclei), *Cntd1*^-/-^ (SKP1 n=26 nuclei and HORMAD1 n=40 nuclei). n ≥3 fetuses were used and data was analyzed using Mean and standard error mean lines are in red. A Mann-Whitney test was utilized to test for statistical significance. P-values are as follows, p<0.05 (*), p<0.001 (**), p<0.0002 (***), p<0.0001 (****).

To further understand the premature acceleration and/or desynapsis in *Cntd1* mutant oocytes, we looked at HORMAD1 accumulation in *Cntd1* oocytes to determine if the oocyte loss we observed in *Cntd1* mutants was also dependent on the synapsis checkpoint, which is monitored by HORMAD1 (Shin et al. 2013; Kogo et al. 2012). We quantified abnormal HORMAD1 in *Cntd1* pachytene-staged oocytes (using a binary system wherein at least one SC had HORMAD1 signal) and found that compared to *Cntd1*^HA/HA^ and *Cntd1* ^HA/-^, *Cntd1*^-/-^ have an increase in the average percentage of oocytes with abnormal HORMAD1 (Fig. 7E-H; *Cntd1*^HA/HA^: 13.13%, *Cntd1* ^HA/-:^ 40.87%, and *Cntd1*^-/-^: 97.33%). These results agree with previous reports that the SCF complex targets HORMAD1 for degradation and removal from synapsed regions of the SC (Guan et al. 2020; Guan et al. 2022). Together, these results indicate that loss of *Cntd1* results in defects in both DSB repair and acceleration / premature desynapsis in prophase I, which may trigger meiotic quality control checkpoints and are likely the causes of significant loss of oocytes (Fig. 3).

### Ablation of CHK2, but not CHK1 rescues oocyte loss in *Cntd1*^-/-^ ovaries

DSB repair and synapsis are carefully monitored throughout prophase I through the action of the checkpoint surveillance kinases ATR and ATM (Martínez-Marchal et al. 2020; Ravindranathan et al. 2022; Huang and Roig 2023). These kinases activate Checkpoint Kinase-1 (CHK1) and Checkpoint Kinase-2 (CHK2), respectively, in response to DNA damage (Rinaldi et al. 2020; Martínez-Marchal et al. 2020; Chen et al. 2012; Bolcun-Filas et al. 2014; Rinaldi et al. 2017; Huang and Roig 2023). Meiotic mutants with an exhausted ovarian reserve phenotype could be rescued by ablating CHK1 and/or CHK2 (Rinaldi et al. 2017; Bolcun-Filas et al. 2014; Martínez-Marchal et al. 2020). Thus, we asked whether the early loss of oocytes in *Cntd1*^-/-^ ovaries could be reversed by loss of either CHK1 and/or CHK2.

Due to the embryonic lethality associated with *Chk1* elimination, we opted for a pharmacological approach using a previously published *in vitro* culture system to determine if oocyte loss in *Cntd1*^-/-^ was CHK1-dependent (Martínez-Marchal et al. 2020; Liu et al. 2000). Post-natal day 5 ovaries were cultured in media containing either DMSO or the CHK1 inhibitor (CHK1i), Rabusertib (LY2603618), which inhibits the autophosphorylation of Serine 296 (S296) and the DNA-damage associated phosphorylation of Serine 345 (S345). This inhibition of phosphorylation prevents CHK1 activation and the subsequent loss of phosphorylation of downstream targets (King et al. 2014) (Fig. 8A). We exposed one ovary per fetus to DMSO and the other to CHK1i. After incubation, we quantified oocytes positive for DDX4, a germ-cell marker in oocytes from both ovaries, as previously published (Martínez-Marchal et al. 2020). CHK1i exposure resulted in no change in the number of oocytes in *Cntd1*^HA/HA^ ovaries exposed to DMSO or CHK1i (Fig. 8B, C, H; mean oocytes ± S.E.M; 992.5 ± 437.5 and 1288 ± 302.5, p=0.2732, respectively), and nor did those from *Cntd1*^HA/-^ ovaries exposed to DMSO or CHK1i (Fig. 8D, E, H; 771.7 ± 430.1 and 510 ± 237.1, respectievely, p=0.3362). Similarly, treatment of *Cntd1*^-/-^ ovaries with DMSO or CHK1i produced no statistically significant change in the number of oocytes (Fig. 8F, G, H; 76.25 ± 32.87 and 320 ± 84.34, respectively, p=0.0979). From these data, we conclude that oocytes in *Cntd1*^-/-^ are not being lost in a CHK1-dependent manner.

**Figure 8.**
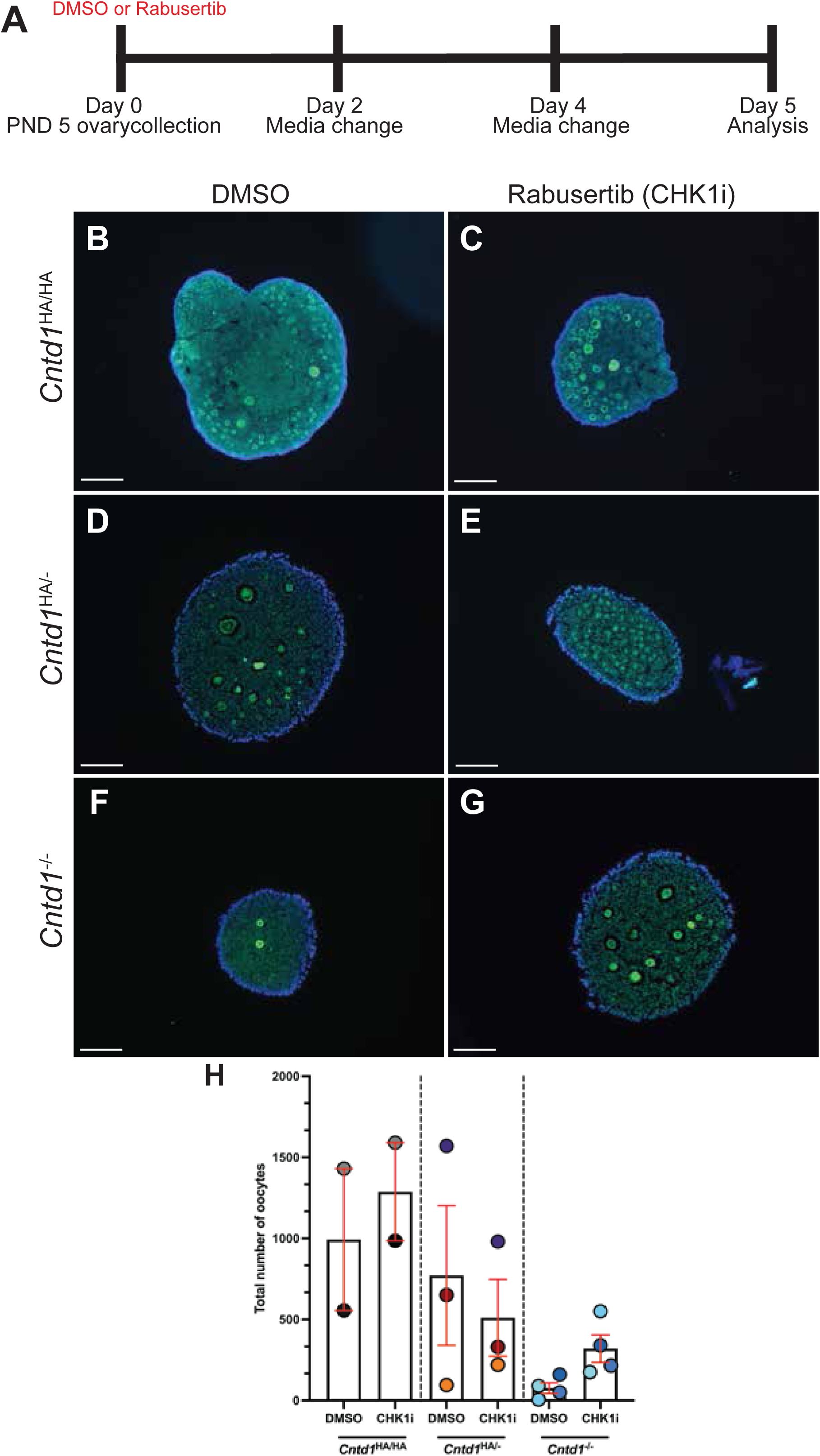
*Chk1* inhibition *in vitro* rescues oocyte loss in post-natal day 5 *Cntd1*^-/-^ ovaries. (A) Culture system outline with Rabusertib for PND5 ovaries. (B-G) Histological sections (5µm) of post-natal day 5 immuno-stained ovaries with DDX4 (green) and DAPI (blue) of *Cntd1*^HA/HA^, *Cntd1*^HA/-^, and *Cntd1*^-/-^. (H) Quantification (mean oocyte number ± SEM) of all DDX4 positive oocytes from *Cntd1*^HA/HA^, *Cntd1*^HA/-^, and *Cntd1*^-/-^ ovaries. Oocytes from n≥2 ovaries were quantified. Scale bar is equal to 100 µm. Mean and standard error mean lines are in red. Significance was determined using a Wilcoxon test. P-values are as follows: p<0.05 (*), p<0.001 (**), p<0.0002 (***), p<0.0001 (****).

Due to the aberrant repair in *Cntd1* mutant oocytes, we next determined if CHK2 was being activated (Fig. 4, 5). To do so, we bred our *Cntd1* mutant line with *Chk2* mutants to generate double homozygous mutant animals (*Chk2*^-/-^;*Cntd1*^-/-^) and associated control genotypes. We analyzed follicle composition from females at PND 28 in compound mutants for *Cntd1* and *Chk2*, including variations of genotypes from fully wildtype females to full double homozygous mutant females (including *Chk2*^+/+^;*Cntd1*^+/+^, *Chk2*^+/+^;*Cntd1*^-/-^, *Chk2*^+/-^; *Cntd1*^+/-^, *Chk2*^-/-^;*Cntd1*^+/+^, and *Chk2*^-/-^;*Cntd1*^-/-^) (Fig. 9A-J). PND28 was selected because this is the age at which we begin to see primordial follicle loss in the *Cntd1* homozygous mutant females (Fig. 3L). If CHK2 is essential for oocyte elimination in the same pathway as CNTD1, we would expect to see a rescue of primordial follicles in the absence of CNTD1, if not all follicle types.

**Figure 9.**
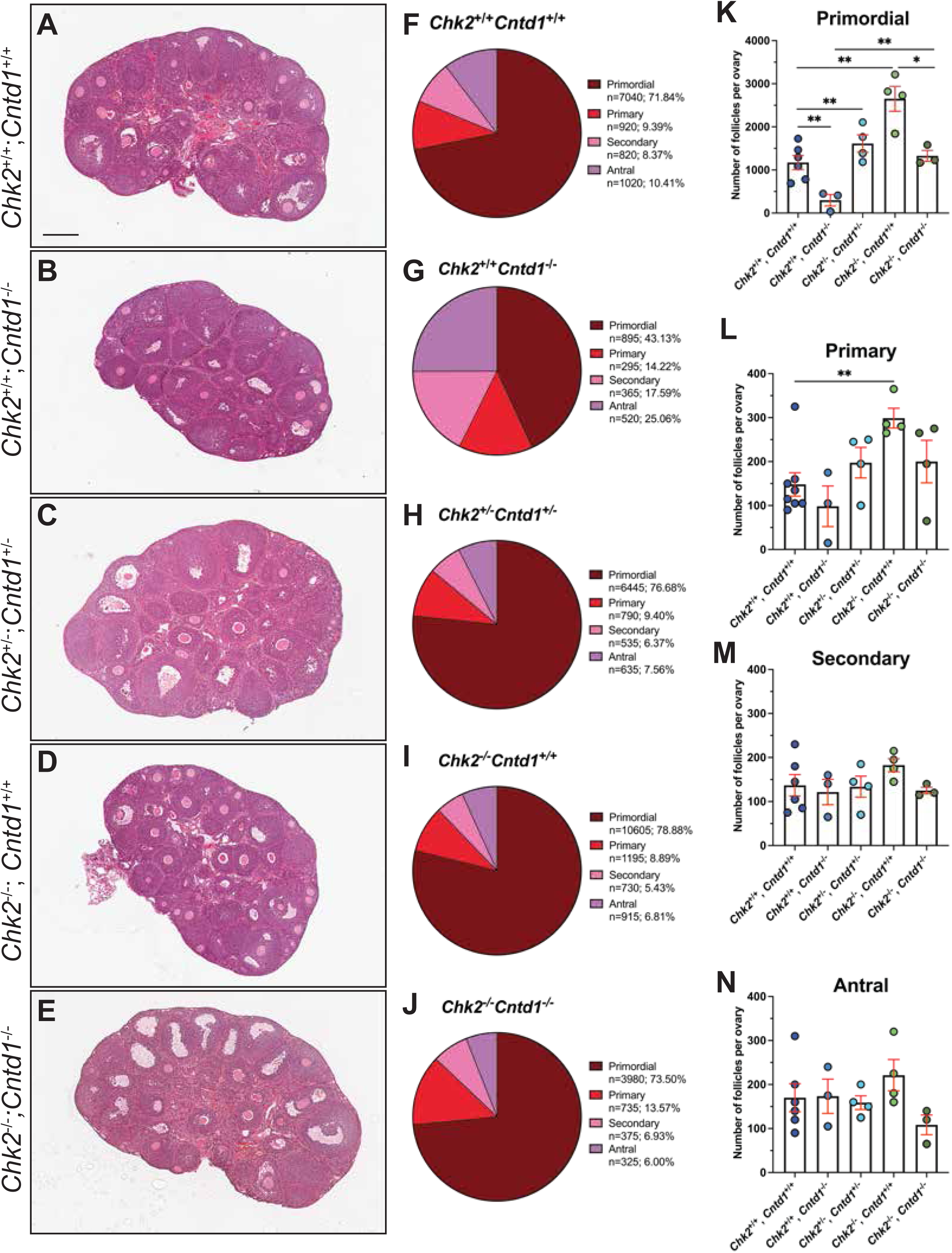
Ablation of *Chk2* results in primordial follicle resuce in *Cntd1*^-/-^ oocytes. (A-E) Histological sections of ovaries from post-natal day (PND) 28 female mice. Paraffin-embedded tissues were sectioned at 5 μm and stained with Hematoxylin and Eosin. Scale bars are equal to 200μm. Composition of primordial (dark red), primary (red), secondary (light pink, and antral (purple) follicle populations for each genotype visualized as a percentage value for (F) *Chk2*^+/+^;*Cntd1*^+/+^, (G) *Chk2*^+/+^;*Cntd1*^-/-^, (H) *Chk2*^+/-^; (I) *Cntd1*^+/-^; *Chk2*^-/-^;*Cntd1*^+/+^, and (J) *Chk2*^-/-^ ;*Cntd1*^-/-^ ovaries. N represents total number of follicles for the respective follicle type with percentage value, calculated as the total number of follicle type divided by the total number of follicles for all follicle types. Quantification (average follicle number ± SEM) of primordial (K), primary (L), secondary (M), and (N) antral follicles. Follicles from n≥3 ovaries were quantified per genotype. Mean and standard error mean lines are in red. Significance was determined using an unpaired t-test with Welch’s t-test. P-values are as follows: p<0.05 (*), p<0.001 (**), p<0.0002 (***), p<0.0001 (****).

The number of primordial follicles was significantly reduced between *Chk2*^+/+^;*Cntd1*^+/+^ and *Chk2*^+/+^;*Cntd1*^-/-^, which was expected due the drastic reduction we observed in *Cntd1*^-/-^ ovaries (Fig. 9K; mean number of follicles ± S.E.M; 1173 ± 163.3, 298.3 ± 132.2, respectively, p=0.0048). There was no difference in the quantity of primordial follicles between *Chk2*^+/+^;*Cntd1*^+/+^ and *Chk2*^+/-^;*Cntd1*^+/-^ (Fig. 9K; 1611 ± 203.7, p=0.1714). Consistent with prior reports of *Chk2*^-/-^ ovaries, we observed an increase in primordial and primary follicles between *Chk2*^+/+^;*Cntd1*^+/+^ and *Chk2*^-/-^;*Cntd1*^+/+^ (Fig. 9K, L; primordial: 2651 ± 290.2, p=0.0071. Primary: 148.1 ± 26.66 and 298.8 ± 22.49, p=0.0018, respectively) (Bolcun-Filas et al. 2014). When we compared the number of primordial follicles between *Chk2*^+/+^;*Cntd1*^-/-^ and *Chk2*^-/-^;*Cntd1*^-/-^, we found that genetically ablating *Chk2* in the absence of *Cntd1* leads to a rescue in the number of oocytes in double homozygous females (Fig. 9K; 1327 ± 127.8). By comparison, we found that the quantity of primordial follicles in *Chk2*^-/-^;*Cntd1*^-/-^ ovaries were reduced compared to *Chk2*^-/-^ ;*Cntd1*^+/+^ (Fig. 9K, p=0.0050). No changes were observed between secondary and antral follicle types in all of the genotypes (Fig. 9M, N). Together, these data demonstrate that *Cntd1*^-/-^ oocytes are being lost through the CHK2 checkpoint pathway.

## DISCUSSION

Defects in crossover regulation and maturation in prophase I lead to the absence of chiasmata and subsequent mis-segregation of homologous chromosomes at the end of meiosis I, resulting in aneuploidy of the gametes (Hassold and Hunt, 2001; Hunt and Hassold, 2008). Such non-disjunction events are particularly prevalent in the first meiotic division in human oocytes compared to spermatocytes (Hassold and Hunt, 2001; Hunt and Hassold, 2008; Nagaoka et al., 2012). This sexual dimorphism can be traced in part back to defects in establishing sites of crossovers during pachynema of meiosis I. Cytological studies on human oocytes revealed that exchangeless chromosomes—synapsed homologous chromosomes lacking a crossover site—and crossovers placed at unfavorable locations are common and account for this high rate of errors (Hassold and Hunt, 2001; Hassold et al., 2021; Nagaoka et al., 2012), and that this correlates with a high level of heterogeneity in the maturation of crossovers in human fetal ovaries (Lenzi et al, 2004). Given these data, studies unraveling the molecular dynamics of crossover regulation in mammalian oocytes are crucial. However, the technical challenges associated with isolating oocytes at early developmental timepoints have hindered such studies in females, with male meiosis being considerably easier to investigate. The current study was aimed at understanding the function of one key regulator of crossover selection, CNTD1, in mammalian oogenesis. We sought to understand the role of CNTD1 in the regulation of crossover designation compared to analogous events in males to better understand the high rate of crossover failure in oocytes compared to spermatocytes.

### Cyclin independent form of CNTD1 is expressed in both mammalian spermatocytes and oocytes

The observation that CNTD1 exists solely as a short form in the testes of male mice was unexpected, but provided a unique mechanism of action for mammalian CNTD1 that is distinct from its worm ortholog, COSA-1 (Gray et al., 2020). Studies presented here confirm this finding and further demonstrate that the short form is the sole form of CNTD1 functioning in murine meiosis in both oocytes and spermatocytes. The current study and previous exploration of CNTD1 in males provides strong evidence for the predominance of CNTD1^SF^ and a cyclin-independent function for CNTD1: (1) an antibody against HA (in *Cntd1-HA* expressing mice) reveals the presence of a single band for CNTD1 in mouse testis and ovary that migrates more quickly than the predicted “full length” CNTD1, thus indicating a smaller sized protein (Fig. 2); (2) our observation of CNTD1^SF^ predominance in mouse testis is confirmed by studies of Bondarieva *et al,* who show a single CNTD1 band that is indicative of short form size using a custom antibody against CNTD1 (Bondarieva et al., 2020), and our unpublished studies with this same custom antibody used on mouse testis protein extracts confirm a similarly smaller-sized protein (Gray and Cohen, unpublished); (3) previous yeast two-hybrid analysis demonstrates that CNTD1^SF^ fails to interact with any CDK, while “full-length” CNTD1 can interact with several CDKs (Gray et al., 2020); and (4) IP-WB as well as IP-MS from mammalian testis lysate using anti-HA antibodies for IP and anti-CDK2 antibodies for WB fail to show an interaction between CNTD1 and CDK2 (Bondarieva et al., 2020; Gray et al., 2020). Thus, mammalian CNTD1 exists only as a short form that lacks a key cyclin-homology domain encoded by exons 1 and 2 of the mouse gene. *In silico* analysis also indicates that transcripts encoding the CNTD1 short form exist in other species as well due to the conservation of a second methionine, including eutherian animals such as human, chimpanzee, koala, dog, and beaver (Gray et al. 2020). In contrast to its predicted role as a cyclin, our previous analysis in male meiosis indicated that endogenous CNTD1 complexes with components of the Replication Factor C (RFC) complex, possibly to recruit the Proliferating cell nuclear antigen (PCNA) clamp to chromosome cores (Gaubitz et al., 2022, 2020). During pachynema, PCNA may then be essential for activating the endonuclease activity of MLH3 to facilitate subsequent crossover formation, although this role for PCNA has only been tested *in vitro* thus far (Cannavo et al., 2020; Kulkarni et al., 2020). Given these data, we speculate that CNTD1 regulates RFC-PCNA localization to sites of MutLγ during crossover designation in both mammalian spermatocytes and oocytes (Gray et al., 2020).

### CNTD1 is essential for the formation of crossovers and synapsis between homologous chromosomes in mammalian oocytes

Repair of all DSBs and full synapsis of homologous chromosomes are essential in oocytes during prophase I. In males, defects in either of these processes is sufficicient to robustly induce quality control checkpoints leading to loss of spermatocytes, while in females, oocytes appear to be somewhat tolerant to such errors. The dual-checkpoint model hypothesizes that both an elevation in unrepaired DSBs and persistent asynapsis beyond zygonema leads to the accumulation of DNA damage response factors and triggers the pachytene checkpoint leading to cell death of meiocytes (Bolcun Filas et al., 2014; Ravindranathan et al., 2022; Rinaldi et al., 2020).

We demonstrate herein that DSB repair events are disrupted in oocytes lacking CNTD1. Specifically, we observe altered RAD51 localization and decreased γH2A.X intensity in pachytene-staged *Cntd1*^-/-^ oocytes (Fig. 4M and Fig. 5D). The persistence of a small number of RAD51 foci in pachynema falls below the published threshold of DSB tolerance (≥10 DSBs) and thus is likely not sufficient alone to trigger the quality control checkpoint (Rinaldi et al. 2017). The reduced γH2A.X intensity may indicate either that DSB repair is not occurring efficiently or that oocytes exhibiting defects in DSB repair may be lost during pachynema and may contribute to loss of oocytes lacking CNTD1. From these data, we speculate that absence of *Cntd1* impacts timely repair of DSBs in prophase I.

The consequence of the delay or disruption of DSB repair in the absence of CNTD1 is associated with the failure to load MutLγ at pachynema, as shown by the complete absence of the MutL homolog, MLH1, a defining feature of class I CO resolution (Fig. 1). The loss of all class I crossovers observed in *Cntd1*^-/-^ females results in dramatic loss of pairing of most homologs at diakinesis, concurrent with loss of 95% of all chiasmata compared to oocytes from *Cntd1^HA^*^/*HA*^ and *Cntd1*^HA/-^ females (Fig. 1). The persistence of 1-2 residual bivalent chromosomes in diakinesis oocytes from *Cntd1*^-/-^ animals is suggestive of an intact class II crossover pathway, presumably catalyzed by the activity of the structure-specific endonuclease MUS81-EME1 (Holloway et al., 2008). The presence of these residual bivalents is also observed in other meiotic crossover mutant mouse strains including *Prr19*, *Rnf212b*, *Mlh1,* and *Mlh3*, further reinforcing the persistence of the class II pathway (Bondarieva et al. 2020; Condezo et al. 2024; Ito et al. 2023; Woods et al. 1999; Kan et al. 2008). Taken together, our data reveal that oocytes lacking *Cntd1* may have inefficient repair of DSBs and that CNTD1 operates exclusively in the class I CO pathway impacting the terminal selection of crossover sites.

Synapsis between homologous chromosoms aids in facilitating the timely repair of DSBs in meiotic prophase I. Defects in establishing and maintenance of synapsis can result in activation of a checkpoint at the end of prophase I (pachytene checkpoint), resulting in the death of oocytes before dictyate arrest (Huang and Roig 2023; Wang and Pepling 2021). Through analysis of fetal oocytes from 15.5dpc through 18.5dpc ovaries, we were able to detect leptotene through diplotene-staged oocytes from all genotypes of *Cntd1* animals (Fig. 6). However, when we performed a systematic progression analysis of prophase I in oocytes from 18.5dpc *Cntd1*^-/-^ oocytes compared to *Cntd1^HA^*^/*HA*^ and *Cntd1^HA^*^/-^ littermate controls, the proportion of pachytene and diplotene oocytes in *Cntd1*^-/-^ is skewed, indicating an acceleration through prophase I, or precocious desynapsis (Fig. 6). Indeed, when we probed putative markers of synaptic defects, we found that *Cntd1* mutant oocytes have both abnormal retention of HORMAD1 and a global decrease of SKP1 intensity along synapsed homologous chromosomes (Fig. 7).

While the meiotic acceleration/asynapsis phenotype reported herein for *Cntd1^-/-^* females is not observed or reported in other crossover regulator mutants such as *Hei10*, *Rnf212*, *Rnf212b*, and *Prr19*, it is reported in *Skp1* conditional knockout oocytes and *Chk2* mutant oocytes (Guan et al. 2022; Martínez-Marchal et al. 2020). SKP1 is a component of the SKP1-Cullin-F-box ubiquitin ligase (SCF) complex, is essential for cell cycle progression, has been shown to be essential for synapsis in mammalian meiocytes, and has been reported to have an independent function as part of the synaptonemal complex in C. *elegans* (Guan et al. 2022; Guan et al. 2020; Blundon et al. 2024). Specifically, *Skp1* conditional knockout meiocytes phenocopy the premature desynapsis/accerlation phenotype we observe in *Cntd1* mutant oocytes. We find that *Cntd1* mutant oocytes have a defect in accumulation of both SKP1 and its target HORMAD1 (Fig. 7), and our lab has reported that CNTD1 in mouse spermatocytes interacts with CDC34, a component of SCF in spermatocytes. From these reports, we speculate that CNTD1 in oocytes may have an independent function outside of CO formation to maintain synapsis in the same pathway as SKP1 during meiotic prophase I.

### Loss of *Cntd1* results in a primary ovarian insufficiency phenotype, which is rescued by genetic ablation of *Chk2*

At the end of prophase I, oocytes are arrested in dictyate until ovulation (Hassold et al., 2021; Morelli and Cohen, 2005; Wang and Pepling, 2021). Dictyate arrested oocytes undergo folliculogenesis beginning in early post-natal life, when the somatic cells of the ovary reorganize into granulosa cells and surround the oocytes to form primordial follicles (Tingen et al., 2009). However, the majority of primordial follicles are lost soon after birth in a process called atresia, in which a large proportion of oocytes undergo apoptosis (Tingen et al., 2009). In humans, atresia leads to the loss of 6 million oocytes, with only roughly 750,000 remaining by 6 months of age (Baker, 1963). While atresia is a normal process in all mammals, there are also examples of certain pathologies that result in excessive loss of oocytes and follicles. The most severe example of this in humans is primordial ovarian insufficiency (POI), in which either an insufficient number of or defective oocytes in human females before the age of 40 can lead to infertility or difficulty with conception (Guo et al., 2017; Huang et al., 2021; Ke et al., 2023; Veitia, 2020). The causes underlying POI are numerous; however, recent Next Generation Sequencing studies have implicated several meiotic genes involved in DSB repair and synapsis, including, *MSH4*, *PRDM9*, *SYCE1,* and *SLX4* (Huang et al., 2021; Ke et al., 2023). Moreover, extensive studies from the Schimenti lab to understand the checkpoint mechanism involved in follicle loss uncovered other meiotic mouse mutants such as *Dmc1*, *Spo11*, *Trip13,* and *Hormad1* which also exhibit a POI-like phenotype, adding more evidence to defective DSB repair and synapsis triggering quality control checkpoints, CHK1 and CHK2 (Bolcun Filas et al., 2014; Di Giacomo et al., 2005; Rinaldi et al., 2020, 2017). More recent studies into the crossover regulators *Rnf212*, *Rnf212b*, *Hei10*, and *Prr19* show that folliculogenesis occurs in these mutants, however the extent of any defects in folliculogenesis remains unknown (Bondarieva et al. 2020; Qiao et al. 2014; Ward et al. 2007; Condezo et al. 2024).

Studies reported herein demonstrate that loss of CNTD1 results in a reduced number of primordial follicles by post-natal day 14 and this reduction continues until adulthood when there is a concurrent loss of all follicle types (Fig. 3D, H, L, P). The early loss of oocytes from birth up until post-natal day 14 is temporally distinct from the two oocyte loss phenotypes reported for other CO regulators. Briefly, loss of *Msh4* or *Msh5,* both of which result in defects from zygonema onwards, results in a similar timing of oocyte loss from birth, albeit much more dramatic and rapid, and necrotic-appearing ovaries by adulthood (Edelmann et al., 1999; Kneitz et al., 2000) (Fig. 10). By contrast, mice lacking *Mlh1* or *Mlh3* have similar meiotic phenotypes to *Cntd1^-/-^* females, showing loss of chiasmata at diakinesis, but no detailed report of alterations to the follicle populations (Woods et al. 1999; Kan et al. 2008; Lipkin et al. 2002).This difference between prophase I function and follicular phenotypic consequence may reflect additional roles of CNTD1 in meiotic cell cycle regulation and/or checkpoint signaling.

**Figure 10.**
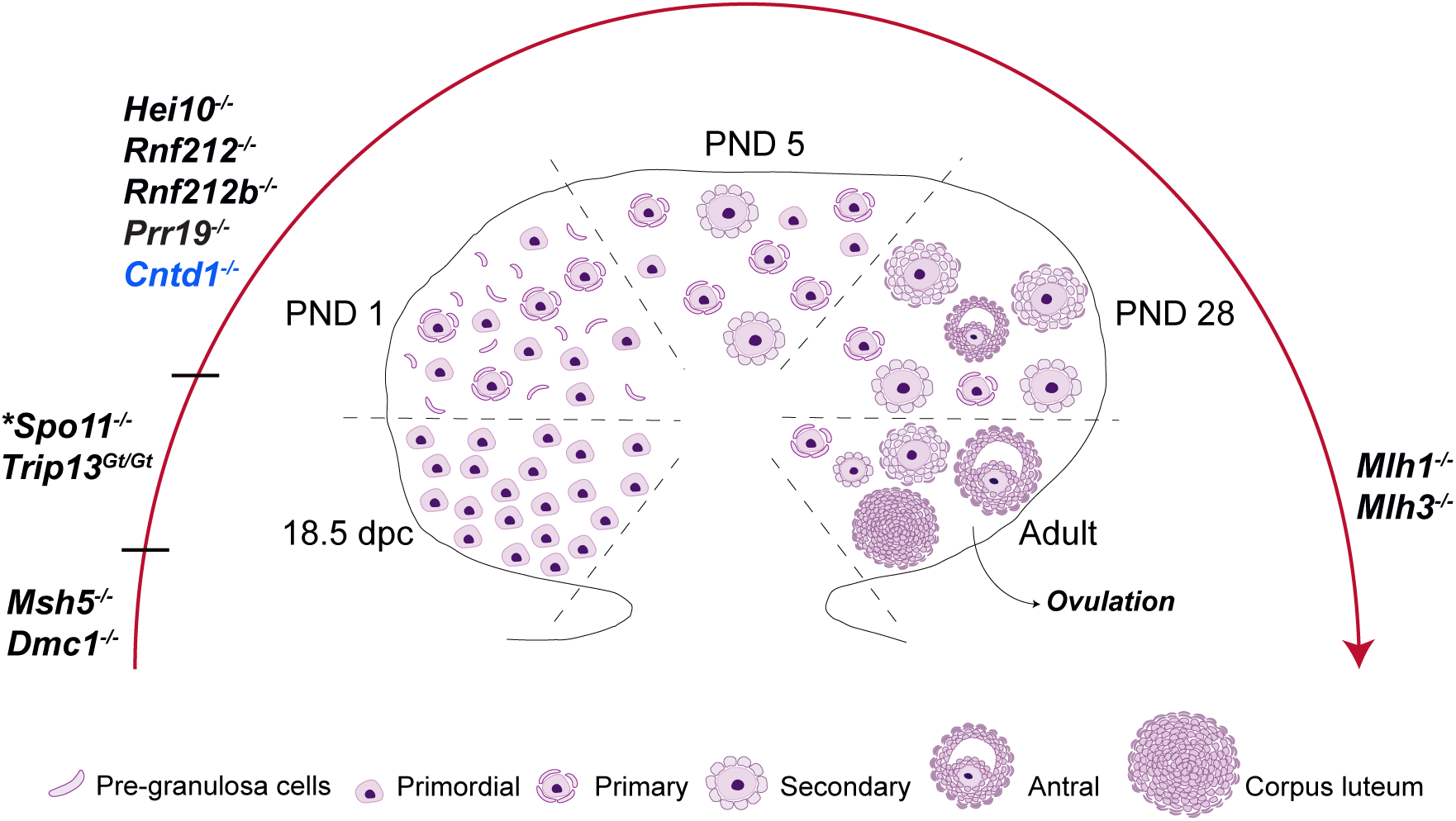
Model of follicle loss in different meiotic mutants. Summary of known ovarian phenotypes for genes important for CO designation and processing through the class I pathway. While some of the phenotypic consequences of these mutations have not been fully described, this summary represents our current understanding of the timing of loss of oocytes in each of the listed mutants. The position of each mutant on the line corresponds to the approximate timepoint of loss of oocytes in that mutant. Genes in blue indicate uncharacterized follicle loss. Loss of oocytes in *Cntd1^-/-^* corresponds roughly to a timepoint after DSB, DNA resection, and CO licensing factors (*Msh5, Dmc1, Spo11, Trip13*) but before CO maturation factors (*Rnf212, Rnf212b, Mlh1, Mlh3, Hei10*). Data extacted from several sources (Bondarieva et al., 2020; Kan et al., 2008; Kneitz et al., 2000; Woods et al., 1999).

The checkpoint kinases *Chk1* and *Chk2* have been shown to be the predominant regulators in oocyte quality control (Martínez-Marchal et al. 2020; Huang and Roig 2023). In the context of meiosis, these kinases are triggered by defects in DNA damage repair and failure of synapsis and subsequent signaling by the upstream effector kinases, ATR and ATM (Rinaldi et al. 2017; Rinaldi et al. 2020; Di Giacomo et al. 2005; Martínez-Marchal et al. 2020). Mouse mutants which bear mutations in DSB repair (*Spo11*) or synapsis (*Trip13* and *Hormad1*) show a similar ovarian phenotype to *Cntd1*^-/-^ animals, with a partial or complete loss of follicles by adulthood (Rinaldi et al. 2017; Bolcun-Filas et al. 2014). Mice deficient in both DSB repair or synapsis genes and *Chk2* show a rescue in oocyte number, showing *Chk2* to be essential for eliminating oocytes containing these deficiencies (Bolcun-Filas et al. 2014; Rinaldi et al. 2020; Rinaldi et al. 2017). Moreover, *Chk2* mutant oocytes phenocopy the synapsis/acceleration defect in prophase I seen in *Cntd1* mutant oocytes (Fig. 6M) (Martínez-Marchal et al. 2020). Ovaries deficient in both *Chk2* and *Cntd1* showed an increase in the number of all follicle types in post-natal day 28 aged animals relative to *Cntd1* single mutant females (Fig. 9E, J, N), whereas follicle numbers in *Cntd1* mutants are not rescued by *in vitro* inhibition of CHK1 activity (Fig. 8H). Together, we speculate that *Cntd1* has a novel role in maintaining oocyte quality before and after the pachytene checkpoint, and its loss results in the apoptosis of oocytes before dicytate arrest and during folliculogenesis.

### Final thoughts and future directions

In mouse spermatocytes, CNTD1 has been proposed to play dual roles during meiosis: in promoting CO formation through an interaction with the RFC/PCNA complex, and in regulating meiotic cell cycle progression through an interaction with the SCF (Gray et al., 2020; Guan et al., 2022, 2020). It is possible that this dual role for mammalian CNTD1, which is unique from its worm ortholog COSA-1, accounts for the earlier loss of follicles in *Cntd1* mutants compared to mutants of proteins similarly essential for Class I COs (*Mlh1/3, Hei10, Rnf212*). Thus, we hypothesize that CNTD1 plays an important role in regulating meiotic cell cycle progression in both sexes, specifically the maintenance of dictyate arrest in oocytes and the prophase I to M-phase progression in spermatocytes. SKP1 has also been shown to be important for maintaining proper synapsis in spermatocytes; *Skp1*cKO spermatocytes driven by a tamoxifen induced Cre show premature desynapsis (Guan et al. 2020). The loss of SKP1 localization to SCs in *Cntd1* mutant oocytes suggests that SKP1 may be targeted to the SCs by CNTD1, where SKP1 acts to maintain synapsis. How SKP1 performs this pro-synapsis role in mice remains unclear; however, recent work in *C. elegans* has shown that worm SKP1 is a core member of the SC, suggesting a possible conserved function (Blundon et al. 2024). It remains to be tested if SKP1 is a component of the SC or if SKP1 is regulating synapsis through the canonical action of the SCF complex in mouse oocytes.

Taken together, the data presented herein reveals two distinct functions of CNTD1 in prophase I in mammalian oocytes. First, CNTD1 ensures that class I COs are established, leading to accurate segregation of chromosomes at the first meiotic division (Fig. 1). Second, CNTD1 participates in regulating the oocyte quality control checkpoint in a cell cycle-depenent manner as seen by the distinct follicle loss phenotype which is distinct from early and late pro-crossover factors and temporally inconsistent with follicle loss through the pachytene checkpoint (Fig. 3 and 9). Future studies are aimed at understanding the specific activity of CNTD1 in regulating crossover sites as well as determining the mechanism by which *Cntd1*^-/-^ oocytes are eliminated, given the defects in synapsis and DSB formation. Further emphasis on female meiosis is likely to yield critical information that elucidates why mammalian oogenesis is so error prone.

## FIGURE LEGENDS

**Supplementary Figure 1. Colocalization analysis of CNTD1 and MLH3.** Representative images of (A-C) *Cntd1*^HA/HA^ (n=12 nuclei) and (E-G) *Cntd1*^HA/-^ (n=19 nuclei) pachytene-staged oocytes (n ≥ 3 fetal ovaries) stained for HA (magenta), MLH3 (green), and SYCP3 (blue). Images are representative of experiments from n ≥ 3 fetuses. Average percentage of colocalization for *Cntd1*^HA/HA^ and *Cntd1*^HA/-^ of MLH3 with HA is 91.53% and 87.34%, respectively. (D and H) Quantification of colocalization. Average percentage of colocalization for *Cntd1*^HA/HA^ and *Cntd1*^HA/-^ of HA with MLH3 is 89.27% and 93.44%, respectively. Mean and standard deviation lines are in red.

**Supplementary Figure 2. *Cntd1*^-/-^ females are sterile.** A 3 month trio breeding assay of *Cntd1*^HA/HA^, *Cntd1*^HA/-^, and *Cntd1*^-/-^ females housed with a *Cntd1*^HA/HA^ male. Total pup number per litter are recorded for each dam genotype. n≥ 3 females were used per genotype. Mean and standard deviation lines are in red.

## AUTHOR CONTRIBUTIONS

**Experimental Design:** Paula E. Cohen, Anna J. Wood, and Ian Wolff

**Experiments:** Anna J. Wood, Rania Ahmed, Leah Simon, and Ian Wolff

**Data analysis and Interpretation:** Anna J. Wood, Rania Ahmed, Ian Wolff, Rachel Bradley, Paula E. Cohen

**Manuscript Preparation:** Anna J. Wood, Ian D. Wolff, and Paula E. Cohen

## ACKNOWLEDGEMENTS

We extend our thanks to Dr. Stephen Gray, Dr. Maria De Las Mercedes Carro, Dr. Tegan Horan, Stephanie Tanis, and Amanda Touey-May for their comments on the manuscript, and to the entire Cohen lab for advice and guidance throughout the course of these studies. We also thank Dr. Carolline Ascenção for her experimental help with co-immunoprecipitations. We are exceptionally grateful to Eileen Shu for outstanding technical assistance and lab leadership. We thank the Cornell Stem Cell and Transgenic Core facility for the creation of the *Cntd1*^HA/HA^ and *Cntd1*^-/-^ mouse lines, Dr. Matthew Thomas from the Cornell Statistical Consulting Unit for his aid in experimental design and data analysis, and the College of Veterinary Medicine Histology Core for tissue embedding and sectioning. Funding for this research was provided by the National Institutes of Health through grants to P.E.C. (HD041012, HD097987), I.D.W. (F32HD106630), and through a scholarship from The Natural Sciences and Engineering Research Council of Canada to RAB (PGSD-577969).

## DELCARATIONS OF INTERESTS

The authors declare no competing interests.

## KEY RESOURCES TABLE

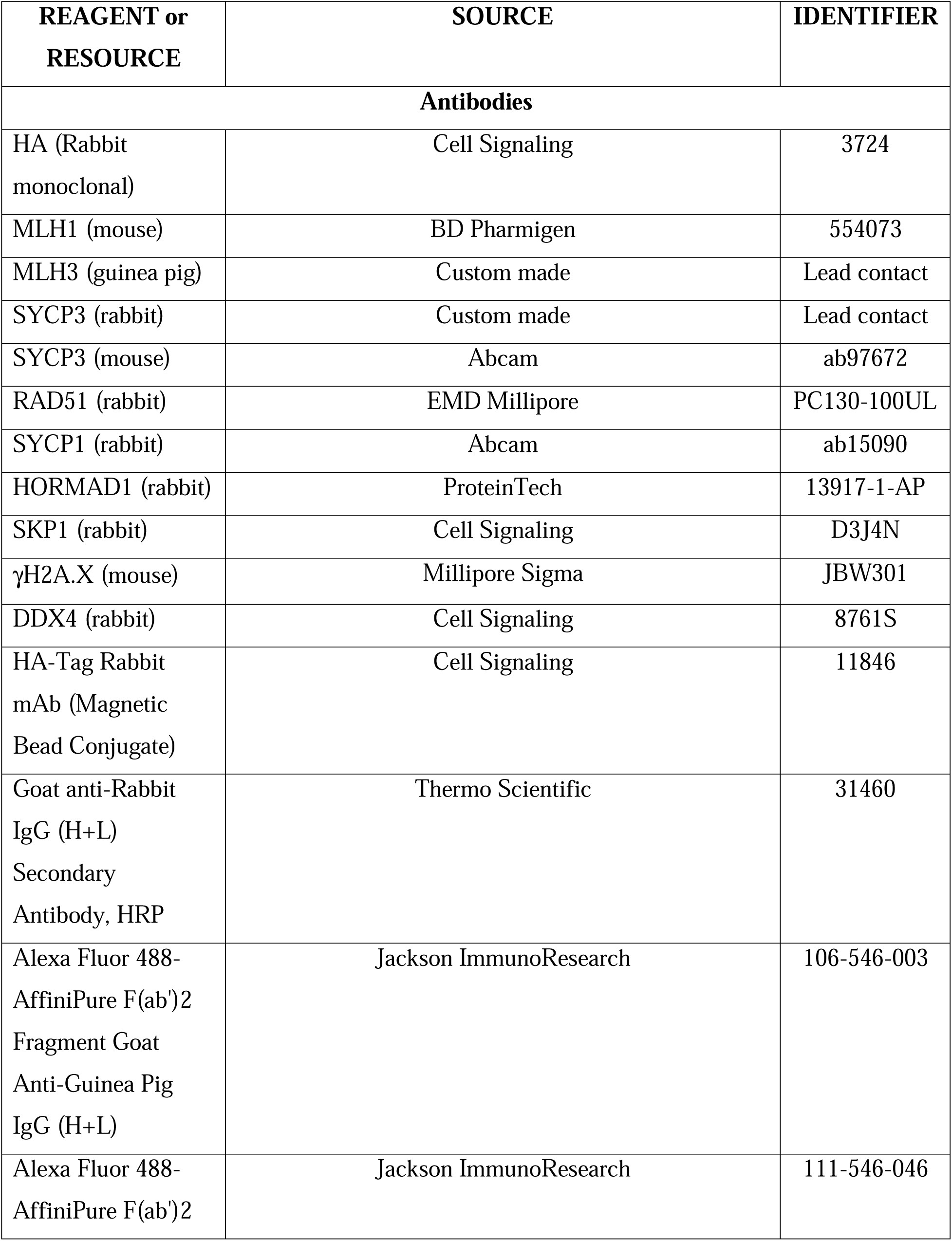
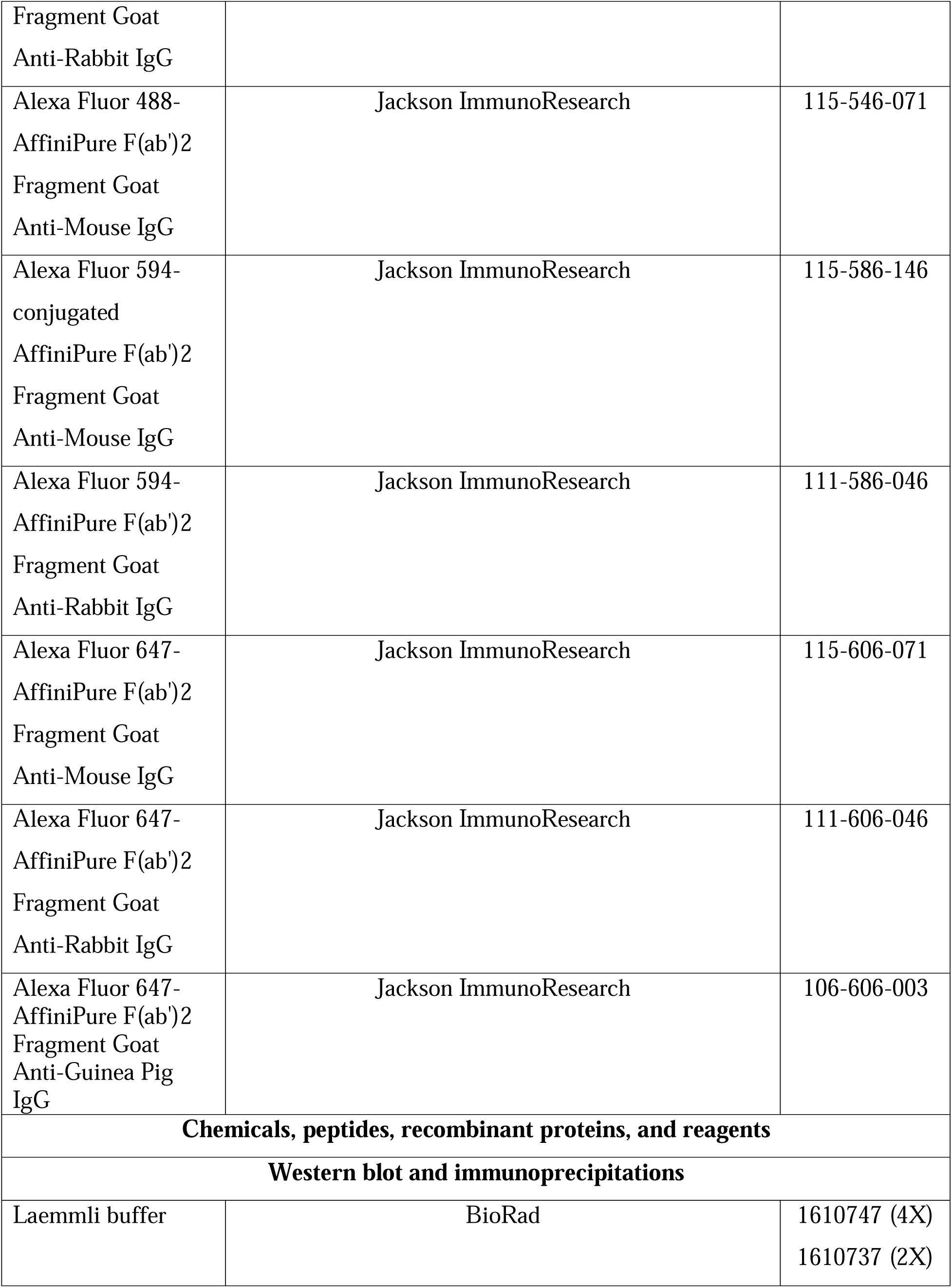
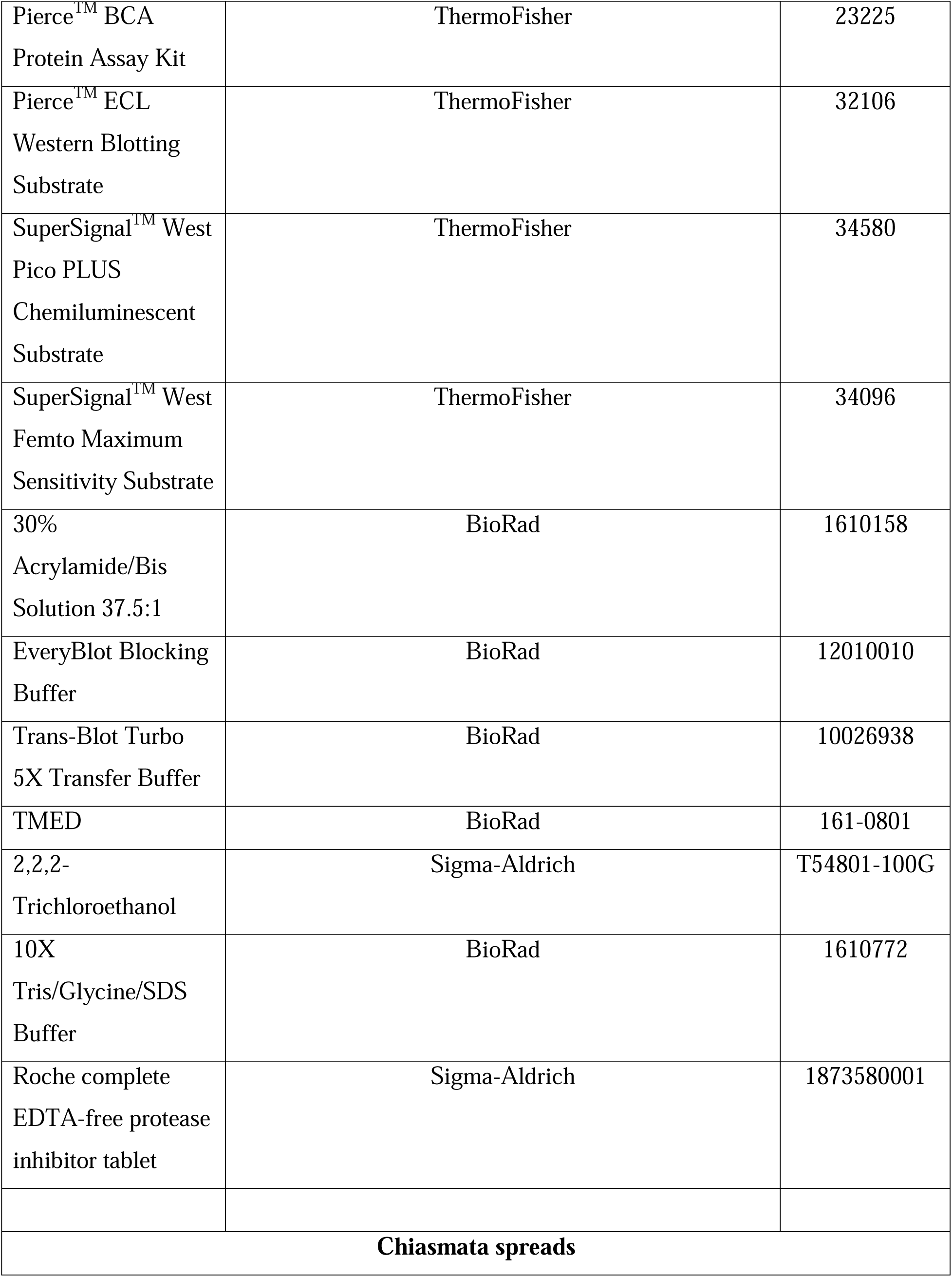
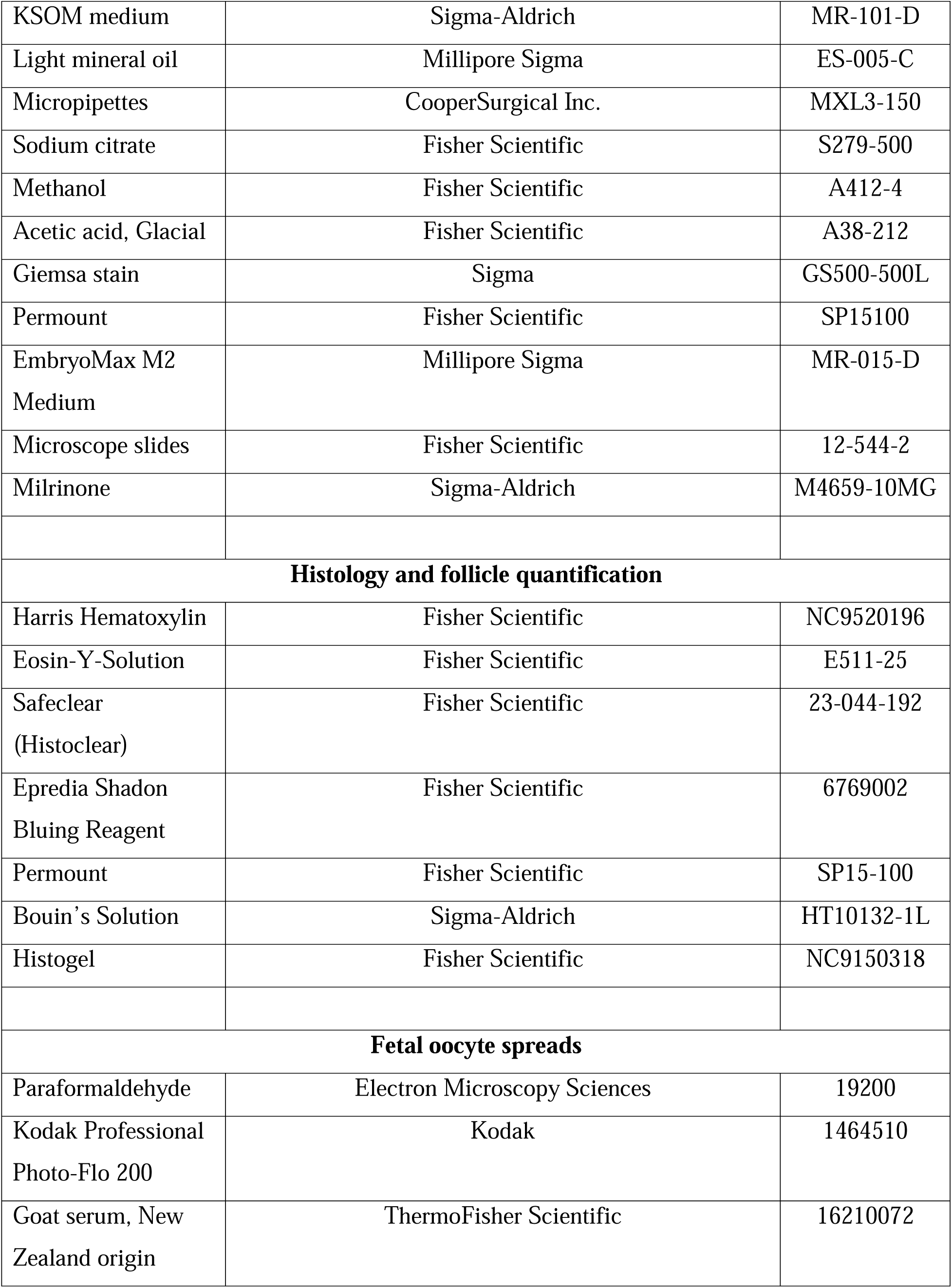
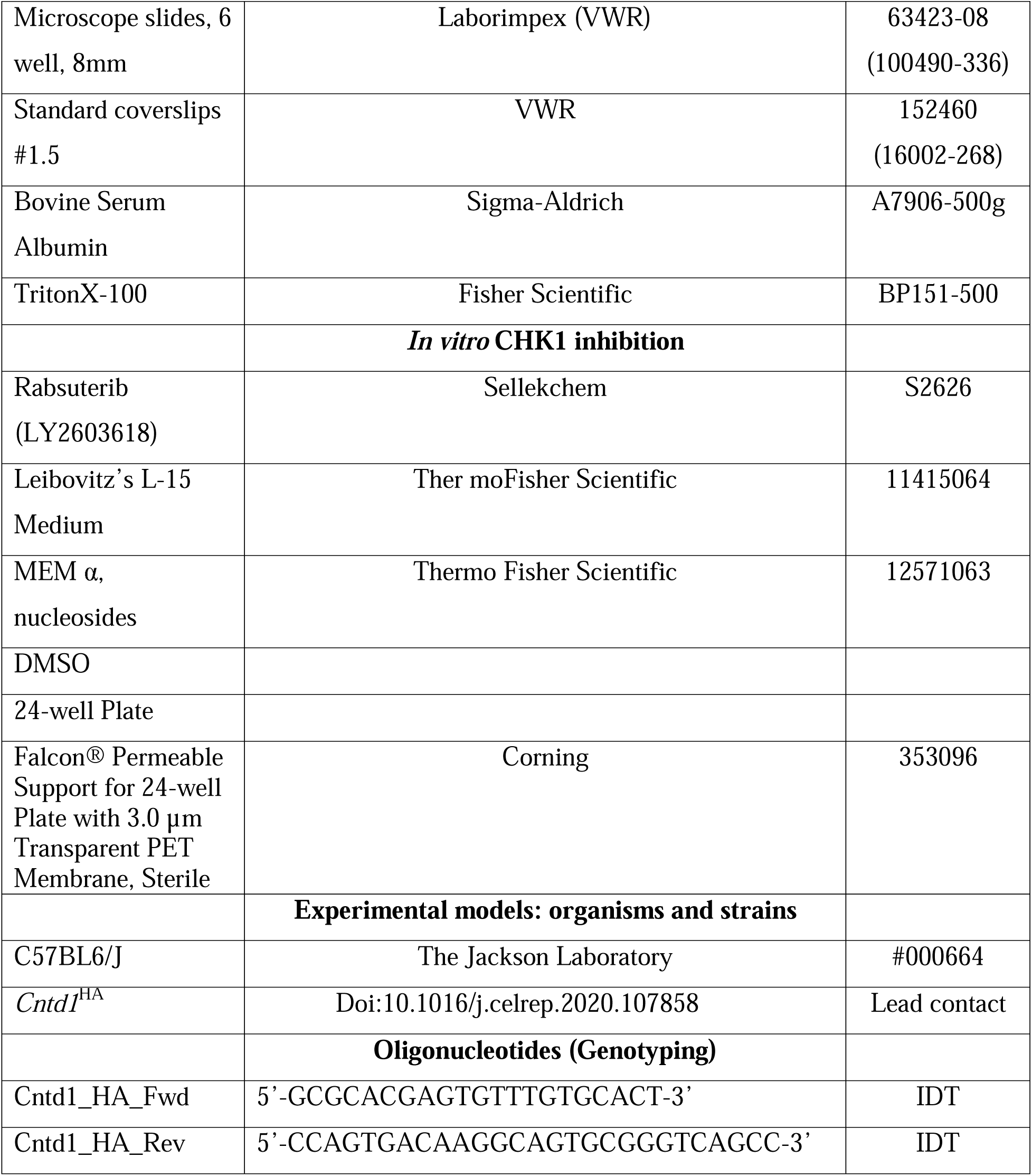

## RESOURCE AVAILABILITY

### Lead contact

Further information and requests for resources and reagents should be directed to and will be fulfilled by the Lead Contact, Paula Cohen

### Materials Availability

Mouse lines generated within this study are available upon request from the Lead Contact, Paula Cohen

### Data and Code availability

This study did not generate any code.

## METHODS

### Mouse strains

All mouse strains were maintained on a C57BL6/J background. *Cntd1*^HA^ and *Cntd1^-/-^* mouse strains were used as reported previously (Gray et al., 2020). All mice were maintained under strictly defined conditions of constant temperature and light:day cycles, with food and water *ad libitum.* Animal handling and procedures were performed following approval by the Cornell Institutional Animal Care and Use Committee, under protocol 2004-0063. All experimental animals were genotyped (Table 1 and Table 2) from ear snips collected and processed with 0.05M NaOH and 1M Tris-HCl pH 7.2. The *Chk2* mutant mice were obtained as a generous gift from Dr. John Schimenti. The *Chk2;Cntd1* compounds were generated initially by mating *Chk2*^+/-^ and Cntd1^+/-^ individuals. Offspring were genotyped (Table 1 and Table 2) and individuals that were *Chk2*^+/-^; *Cntd1*^+/-^ were used for both experiments and maintenance of the line.

### Breeding assay

Two *Cntd1*^HA/HA^, *Cntd1*^HA/-^, or *Cntd1*^-/-^ females and one *Cntd1*^HA/HA^ male were allowed to mate for up to 3 months. Each trio was allowed to produce pups up to three months from the pairing date. Subsequent litter numbers were recorded at weaning.

### Chromosome spreading and immunofluorescence

Oocytes were isolated from 16.5 to 18.5 days post-coitum (dpc), with the day of observing the vaginal plug being 0.5 dpc. Ovaries were dissected into 1X PBS and incubated in hypotonic extraction buffer (HEB) for approximately 20-30 minutes at room temperature, which comprised of Tris-HCl (pH 8.2), 50 mM sucrose, 17 mM trisodium dihydrate, 5 mM EDTA, 0.5mM DTT, and 0.1 mM PMSF. After incubation the ovaries were punctured in 100 mM sucrose and added to slides containing 1% paraformaldehyde with 20% Triton-X. The ovaries were allowed to fix at room temperature for 2 hours in a humid chamber. After 2 hours, slides were allowed to dry at room temperature. Slides were washed in a mixture of 0.4% Photoflo and miliQ-H_2_O once for 5 minutes then allowed to dry completely. For preparation for primary antibody staining, slides were washed with 0.4% Photoflo in 1X PBS once for 10 minutes. This was followed by washing in a Triton wash comprised of 0.5% Triton and 10% Antibody Dilution Buffer (“ADB”: 3g BSA, 90mL 1X PBS, and 20% Triton-X, filter sterilized) in 1X PBS once for 10 minutes followed by blocking in 10% ADB in 1X PBS for 30 minutes at room temperature. Slides were subsequently stained with the antibody of interest (HA 1:500, MLH1 1:50, SYCP3 [mouse] 1:500, MLH3 [guinea pig] 1:500, SYCP3 [rabbit] 1:1000, RAD51 1:500, SYCP1 [rabbit] 1: 5000, HORMAD1 [rabbit] 1:1000, SKP1 [rabbit] 1:50, γH2A.X [mouse] 1:1000) overnight at room temperature in a humid chamber. The following day, slides were washed with 0.4% Photoflo in 1X PBS, Triton wash, and blocked in 1X ADB as stated above. Secondary Alexa-fluor antibodies were diluted 1:2000 in 10X ADB and slides were stained for 2 hours at room temperature. Following secondary antibody staining, slides were washed in the dark in 0.4% Photoflo in 1X PBS (three 5-minute washs) at room temperature, followed by a wash in 0.4% Photoflo in miliQ-H_2_O once for 5 minutes. Slides were allowed to dry and mounted with DAPI plus Antifade (8 ng/mL) and sealed with rubber cement and imaged.

### Quantitation and assessment of chromosome spread staining

For staining intensity analyses of H2A.X and SKP1, all images were taken at the same exposure time and run through a previously published script in ImageJ (Alexander et al. 2023). The ratio of 488 (GFP) divided by DAPI were taken for each cell and plotted in Prism. Normalization and descriptive statistics were recorded and statistical significance was determined using a Mann-Whitney t-test. For analysis of HORMAD1 staining, all images were taken as explained above. To declare “abnormal” HORMAD1 staining, at least one SC had HORMAD1 signal present. Values are recorded as the percentage per fetus.

### Oocyte prophase I sub-staging for progression analysis

The presence of a fork at the end of a long and filamentous synaptnemal complex (SC) as visualized by SYCP3 and no localization of SYCP1, but no bubble on the axis were considered as zygotene-staged. Oocytes with co-localized SYCP1 and SYCP3 along the entire axis length of the SC were categorized as pachytene-staged. The presence of a bubble as marked by the presence of SYCP3, but not SYCP1, and/or a fork with no SYCP1 was considered diplotene-staged. Cells with the complete absence of SYCP1 and compact axes as marked by SYCP3 were also considered as diplotene-staged.

### Ovary Histology and quantification

One ovary from mice at postnatal days 1, 14, and 28, and 8–12-week-old adults were fixed in Bouin’s or Formalin for 4 hours at room temperature on a nutator. Following fixation, ovaries were washed in 70% ethanol. For histological preparation, ovaries were embedded in Histogel before being prepared in a cassette. The Cornell Histopathology core serial sectioned Histogel and paraffin embedded ovaries in 5 micrometer sections. Hematoxylin and Eosin staining were performed as follows: three 5-minute washes in Histoloclear, followed by three 3-minute washes in 95% ethanol, subsequently followed by one 5 minute wash in 80% ethanol and 70% ethanol, with a final wash in 1X PBS for 10 minutes. Staining comprised of one 30-second staining in Hematoxylin followed by two washes in miliQ-H_2_O, followed by two dunks in Bluing reagent and subsequent washes in miliQ-H_2_O. Slides were stained with Eosin for 2 minutes followed by four dunks in miliQ-H_2_O. Slides were then processed through 1 dunk of 70% ethanol and 80% ethanol, and three 3-minute washes in 95% ethanol, finished with three 3-minute Histoclear washes. Slides were dried and mounted with Permount and subsequently imaged. Sections were imaged on Aperio software using an Aperio CS2 Digital Pathology Slide Scanner microscope. The total number of follicles was determined by multiplying the raw counts from every fifth section by 5 to correct for the sections not counted. Follicles were staged as explained previously and follicles with oocytes without a visible nucleus were excluded from quantification to avoid double counting (Sarma et al., 2020).

### Oocyte metaphase spread preparations

Spreads were prepared as previously described in Sun and Cohen (Sun and Cohen, 2013). Briefly, M2 collection media was made comprised of Waymouth’s medium, FBS, 1% Pennicillin-Streptomycin, and 2.5 mg/mL Sodium Pyruvate. A 60 mm plastic petri dish was prepared with 20 μ*L* KSOM droplets and Light Mineral oil covering the droplets. Collection media and the KSOM droplets with light mineral oil were equilibrated overnight in a 37°C incubator (5% CO_2_). Ovaries from unstimulated female mice aged 24-28 days were collected directly into warmed collection media. The ovaries were punctured using 26 gauge needles to release oocytes. Oocytes with a visible zona pellucida and cumulus cells were collected using a 150-striper tip and placed into one of the KSOM droplets. Dissociation of the cumulus cells was achieved by mouth pipetting oocytes through the KSOM droplets until cumulus cells were gone. Oocytes in KSOM and mineral oil were incubated for approximately 7-8 hours until the oocytes entered metaphase I. Oocytes were moved to 20 μL droplets of 1% hypotonic solution (1g sodium citrate dissolved in 100 mL miliQ-H_2_O) for 15 minutes. A glass slide prepared with a China pen or hydrophobic pen in a grid formation were used. 1-2 μL of hypotonic solution was placed onto one square on the glass slide. Oocytes without the zona pellucida were transferred to the drop. Excess hypotonic solution was siphoned off to allow the oocytes to adhere to the slide. One drop of Carnoy’s fixative (3 parts methanol and 1-part glacial acetic acid) was added directly onto the oocytes. The fixative was allowed to disperse, followed by another 1-2 drops of Carnoy’s fixative. The slides were allowed to air dry. This was followed by Giemsa staining (48 mL miliQ-H_2_O and 2 mL Giemsa) in a coplin jar for 3 minutes. The slides were washed 3 x 3 minutes in miliQ-H_2_O. The slides were allowed to dry at room temperature, immediately followed by mounting with Permount and a coverslip. Chiasmata were imaged at 63X.

### Immunoprecipitation of proteins from fetal mouse ovarian cell extracts

Co-immunoprecipitations were performed as reported previously with modifications (Gray et al., 2020). Approximately 40 ovaries per genotype (*Cntd1*^HA/HA^ or C57BL6/J) from 18.5 dpc embryos were isolated and snap frozen on dry ice, then stored at -80°C until use. *Cntd1*^HA/HA^ testes were collected from 10 week old adults, detunicated, and snap frozen on dry ice before storage at -80°C until use. Ovaries or testis were added to 1 mL of cold lysis buffer (50 mM Tris-HCl pH 8.0, 0.2% NP-40, 150 mM NaCl, 5 mM EDTA, 0.1 mg/mL PMSF, Roche complete EDTA-free protease inhibitor tablet) and sonicated with the following parameters: 23% amplitude for 12 seconds, 0.4 seconds on / 0.2 seconds off, and rest on ice for approximately 30 seconds between runs. Sonicated tissues were then spun down at 15,000 x g for 20 minutes at 4°C and the supernatant was collected. 20 μL of lysate was removed for BCA quantification and protein amounts were quantified based on manufacturer protocol. 10 μL of rabbit anti-HA conjugated magnetic beads were added to 500 µL of lysate (250 µg total protein) and incubated overnight at 4°C. Unbound supernatant was removed from beads, and beads were washed 4 times for 5 minutes in 500µL ice cold wash buffer (lysis buffer with 250 mM NaCl). Beads were then resuspended in 100µL of wash buffer and transferred to a fresh tube. Bound proteins were eluted from beads by resuspending in 30µL elution buffer (100 mM Tris-HCl pH 8.0, 1% SDS, and 10 mM DTT) then incubating at 65°C for 15 minutes. Elutant was then collected, added to 10 μL 4X Laemmli sample buffer and boiled for 5 minutes at 95°C. Input gel samples were taken before addition of HA beads: 30 μL of lysate was added to 10 μL 4X Laemmli sample buffer (BioRad) and boiled at 95°C for 5 minutes.

### SDS-PAGE and Western Blotting

Protein samples were separated by SDS-PAGE on gels varying in percentage from 6%–14% and transferred to methanol activated PVDF membranes using a Biorad Mini Trans-Blot Cell. Membranes were blocked by incubating in EveryBlot Blocking Buffer at room temperature while rotating at 60 rpm for 10 minutes. Membranes were then incubated overnight in primary antibody diluted in EveryBlot Blocking Buffer at 4°C. Membranes were washed three times in 0.1% TBST and subsequently incubated with secondary antibody for 2 hours at room temperature, followed by three 0.1% TBST washes. Membranes were developed using various ECL reagents and imaged using a Biorad ChemiDoc imager. Antibodies used in this study are described in the Key Resources Table.

### Neonatal ovarian organ cultures in the presence of CHK1 inhibitor

Post-natal day 5 ovaries were dissected and cultured as published previously (Martínez-Marchal et al. 2020; Morgan et al. 2015). Briefly, tail snips were collected at the time of dissection for genotyping with *Cntd1*_HA primers. Ovaries were collected from post-natal day 5 females in dissection media (L15 media with 3mg/mL of BSA). Ovaries were either cultured with 5µM of DMSO or CHK1 inhibitor in α-MEM with 3mg/mL BSA in a 24-well plate with a well insert. Ovaries were cultured in an incubator at 37°C supplied with 5% CO_2_ for five days, with media changes every 48 hours days. Ovaries were then fixed in formalin for 1 hour at room temperature and outsourced to the Cornell Histology Core for subsequent sectioning at 5µm per section. Sections were then deparaffinized with Histoclear and dehydrated through ethanol dilutions. Sections were processed for immunostaining with antigen retrieval in boiling Sodium Citrate Buffer (Tri-sodium citrate 10mM, pH 6.0) for 30 minutes. Immunofluorescence was performed using rabbit anti-DDX4 (Cell Signaling) at 1:1000 in 1X ADB at room temperature for 1 hour and anti-rabbit Alexa Fluor 488 at 1:500 in 1X ADB for 1 hour at room temperature. Slides were mounted with DAPI plus antifade and imaged utilizing a Zeiss Axiophot Z1 microscope at 10X magnification. Every 5^th^ section was analyzed and only DDX4-positive oocytes with a visible nucleus were quantified. The total number of oocytes was multiplied by 5.

### Image acquisition

Imaging was performed using a Zeiss Axiophot Z1 microscope attached to a cooled charge-coupled device (CCD) Black and White Camera (Zeiss McM). Images were captured and pseudo-colored using ZEN 3.0 software (Carl Zeiss AG, Oberkochen. Germany). The brightness and contrast of images were adjusted using ImageJ (Schindelin et al., 2012) (National Institutes of Health, USA).

## QUANTIFICATION AND STATISTICAL ANALYSIS

Statistical analyses were performed using GraphPad Prism version 9.00 for Macintosh and Microsoft Excel. Specific analyses are described within the text and the corresponding figures. All datasets were analyzed for normality using a Shapiro-Wilk test. Alpha value was established at 0.05 for all statistical tests. The statistical tests utilized are included in the figure legends. All statistical analysis was monitored by the Cornell Statistical Consulting Unit.

