## Supplementary figures and images for "CNTD1 plays crucial roles in prophase I progression and crossover designation during female meiosis and is critical for establishing the ovarian reserve"

### Supplemental Figure 1

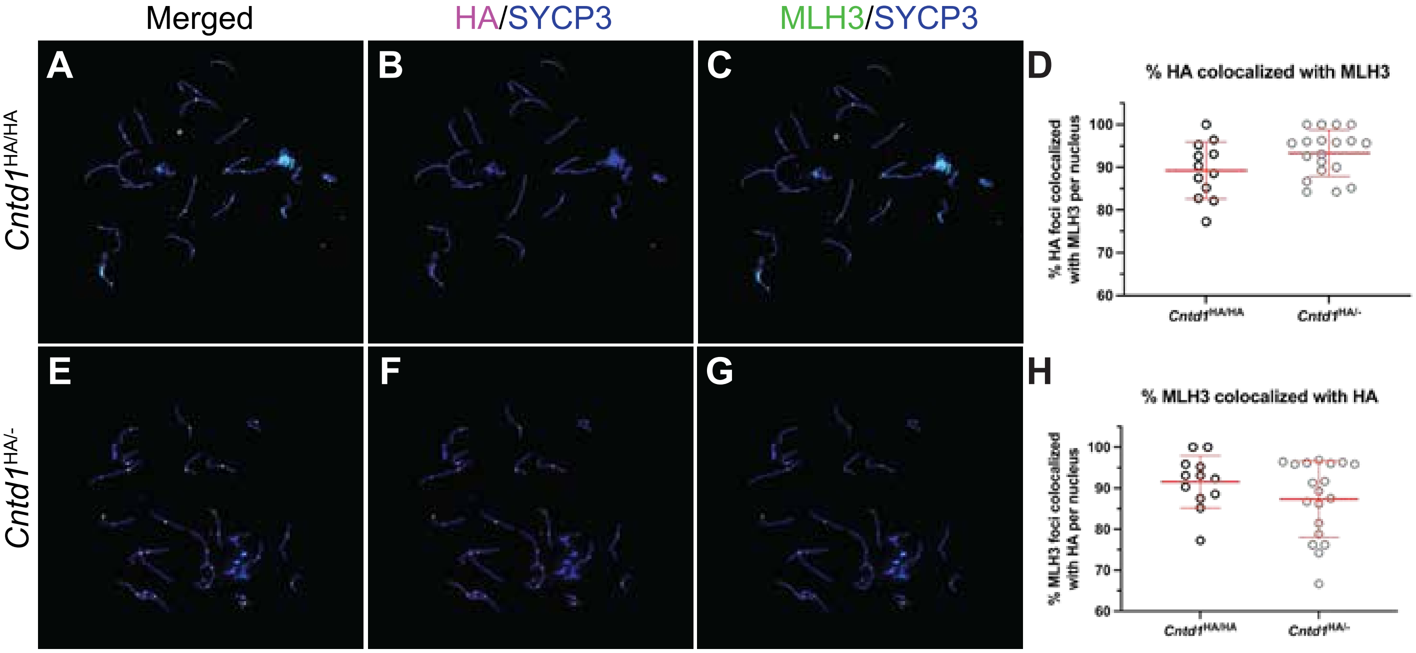

### Supplemental Figure 2

A

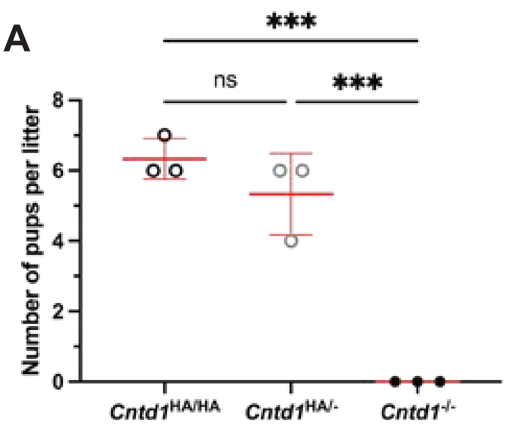
